# A method for three-dimensional single-cell chronic electrophysiology from developing brain organoids

**DOI:** 10.1101/2021.06.22.449502

**Authors:** Paul Le Floch, Qiang Li, Ren Liu, Kazi Tasnim, Siyuan Zhao, Zuwan Lin, Han Jiang, Jia Liu

## Abstract

Human induced pluripotent stem cell-derived brain organoids have shown great potential for studies of human brain development and neurological disorders. However, quantifying the evolution and development of electrical functions in brain organoids is currently limited by measurement techniques that cannot provide long-term stable three-dimensional (3D) bioelectrical interfaces with brain organoids during development. Here, we report a cyborg brain organoid platform, in which 2D progenitor or stem cell sheets can fold “tissue-like” stretchable mesh nanoelectronics through organogenesis, distributing stretchable electrode arrays across 3D organoids. The tissue-wide integrated stretchable electrode arrays show no interruption to neuronal differentiation, adapt to the volume and morphological changes during organogenesis, and provide long-term stable electrical contacts with neurons within brain organoids during development. The seamless and non-invasive coupling of electrodes to neurons enables a 6-month continuous recording of the same brain organoids and captures the emergence of single-cell action potentials from early-stage brain organoid development.

## Introduction

The ability to record tissue-wide, millisecond-timescale single-cell electrophysiology over the time course of human brain development is important to understand the emergence of orchestrated neuronal activities^1^ and elucidate origins of neurodevelopmental diseases^2,3^. This ability has not yet been achieved due to the inaccessibility of the human brain at the early developmental stages. Recent breakthroughs in the development of human induced pluripotent stem cells (hiPSCs) have introduced techniques to grow brain organoids from *in vitro* cultured stem cells that can proliferate, differentiate and self-assemble^4–6^ into three-dimensional (3D) tissues, resembling the cellular architecture, diversity, and electrophysiology of the human brain at early stages^7,8^. Brain organoids thus provide a reliable and easily accessible platform to study human brain development and neurodevelopmental diseases^9–12^, bridging the gap between animal research and human clinical study.

However, long-term stable recording of single-cell electrophysiology in developing brain organoids is still a challenge. The recording technology not only needs to form minimally-invasive and long-term stable electrical interfaces with individual neurons 3D distributed across brain organoids, but also needs to accommodate the rapid volume change occurring during the organoid organogenesis and cortical expansion. Optical imaging coupled with fluorescence dyes^13^ or calcium indicators^14^ has been used to visualize the neuron activities in 3D. They, however, are limited by the temporal resolution, penetration depth, and long-term signal stability. Electrical measurement techniques such as 2D microelectrode arrays (MEA)^15,16^ and patch-clamp^17,18^ have been applied to measure the functional development of brain organoids, but they can only capture the activities from the bottom surface of brain organoids^19–21^ or assay one cell at a time with cell membrane disruption. The recent development of 3D bioelectronics enables 3D interfaces with brain organoids^1,19,21–25^. However, they can only contact organoids at the surface by flexible electronics or penetrate organoids invasively by rigid probes, which cannot further accommodate volume and morphological changes of brain organoids during development. To date, it is still a challenge to non-invasively probe neuron activity at single-cell-single-spike spatiotemporal resolution across the 3D volume of brain organoids and over the time course of development. This constraint prevents further understanding of the functional development in brain organoids and standardizing culture conditions and protocols for brain organoid generation by quantifying their electrical functions.

To address this challenge, we developed the cyborg organoid platform by integrating “tissue-like” stretchable mesh nanoelectronics, which possesses tissue-level mechanical properties and subcellular-scale features, with 2D stem cell sheets. Leveraging the 2D-to-3D reconfiguration during organoid organogenesis, 2D stem cell sheets fold and embed stretchable mesh nanoelectronics with electrodes throughout the entire 3D organoid for continuous recording^26^. Here, we design stretchable mesh nanoelectronics to build cyborg human brain organoids. Using the 3D embedded stretchable electrodes, we achieved reliable long-term electrical recording of the same brain organoids at single-cell, millisecond spatiotemporal resolution for 6 months, revealing the evolution of the tissue-wide single-cell electrophysiology over brain organoid development. Applying this technology to brain organoids at early developmental stages, we traced the gradually emerged single-cell action potentials and network activities.

## Results

### Design of stretchable mesh nanoelectronics

In general, there are two types of protocols to build human brain organoids: i) hiPSCs are first differentiated into neuron progenitors and neurons before self-organizing into 3D structures through organogenesis^27,28^; and ii) hiPSCs first self-organize into spheroids through cell-cell and cell-extracellular matrix (ECM) interactions and then are differentiated into neurons^7,8^. To examine the general feasibility, we built two different types of cyborg brain organoids according to these two types of protocols (Fig. 1a). In type A brain organoids, the hiPSCs (Stage I) were differentiated into neurons by 2D culture for 4 months. After confirming the spontaneous action potentials from the hiPSC-derived neurons by a 2D microelectrode array, cells were dissociated into single cells, integrated with the stretchable mesh nanoelectronics, and induced to self-organize into 3D structures through organogenesis (Stage II-A). In type B brain organoids, hiPSCs were cultured on the stretchable mesh nanoelectronics/Matrigel hybrid structure to form a 3D structure and then induced for neuronal differentiation (Stage II-B). For both methods, the 2D-to-3D reconfiguration folds the 2D cell plate/nanoelectronics hybrids into 3D structures with stretchable mesh electrodes seamlessly distributed across 3D brain organoids. Ultimately, the 3D embedded electrodes (Stage III) are connected with an amplification and data acquisition system to continuously monitor the evolution of electrical signals from neural progenitors and neurons.

**Figure 1.**
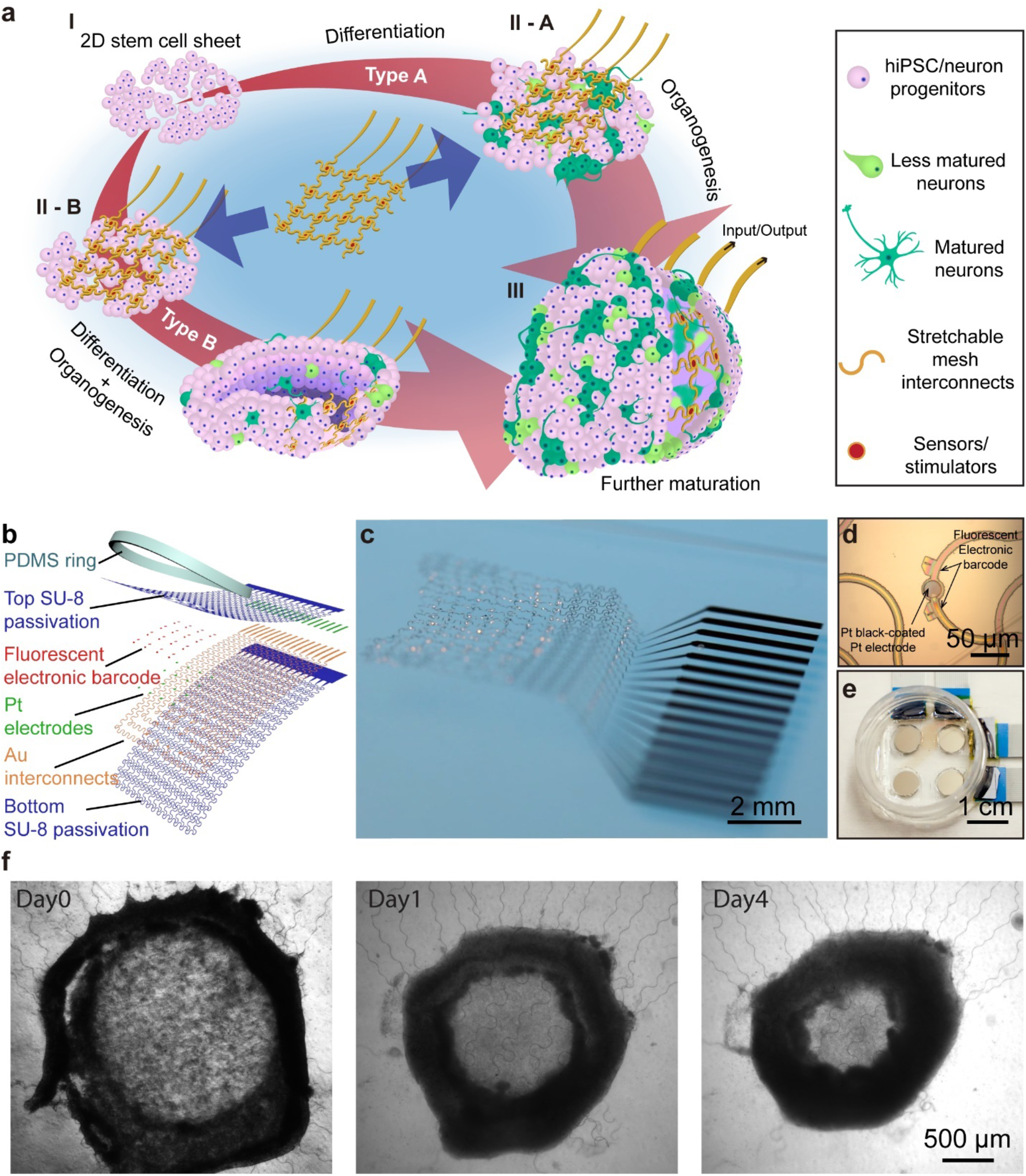
Cyborg brain organoids. **a**, Schematics illustrate the stepwise integration of stretchable mesh nanoelectronics into brain organoids through organogenesis. (I) Human induced pluripotent stem cells (hiPSCs) were seeded with Matrigel. (II) Lamination of stretchable mesh nanoelectronics onto the two-dimensional (2D) cell sheet after neuronal differentiation into neural progenitors (II-A) or lamination of stretchable mesh nanoelectronics onto the hiPSCs before neuronal differentiation (II-B). (III) The organogenetic 2D-to-3D self-organization folds the 2D cell sheet/nanoelectronics hybrid into a 3D structure. 3D embedded sensors are connected to external recording electronics to keep monitoring the electrophysiology of hiPSC-derived neurons and neural progenitors. **b**, Exploded view of the stretchable mesh nanoelectronics design consisting of (from top to bottom) an 800-nm-thick top SU-8 encapsulation layer, a 50-nm-thick platinum (Pt) electrode layer electroplated with Pt black, a 40-nm-thick gold (Au) interconnects layer, and an 800-nm-thick bottom SU-8 encapsulation layer. The serpentine layout of interconnects is designed to enable stretchability. A polydimethylsiloxane (PDMS) ring is bonded around the device as a chamber to define the size and initial cell number in the seeded hiPSC sheet. **c**, Optical photograph of stretchable mesh nanoelectronics released from substrate and floating in the saline solution. **d**, Optical bright-field (BF) microscopic image of stretchable mesh nanoelectronics before released from the fabrication substrate shows a single Pt electrode coated with Pt black. **e**, Optical photograph of a 2×2 devices well, with a single culture chamber for 4 cyborg brain organoids cultured simultaneously. **f**, Optical phase image of stem cells integrated with stretchable mesh nanoelectronics from Day 0 to Day 4 showing that the 2D cell sheet with embedded stretchable mesh nanoelectronics self-folded into a 3D cyborg brain organoid.

To successfully integrate stretchable mesh nanoelectronics with brain organoids, we designed the device structure to accommodate the structural and mechanical properties of developing brain organoids. First, the mesh exploits a serpentine layout with an overall filling ratio of less than 7% and an in-plane stretchability of up to 30% (Fig. 1b-c, Supplementary Fig. 1-3, Methods)^29^. The structure also allows compression and folding through buckling of ribbons in the mesh network^30^. Together, these multiscale deformations can accommodate the compression, folding, and expansion during brain organoid organogenesis^31^. Second, brain organoids may be softer than cardiac organoids^26^, given that brain tissues (elastic modulus of a few kPa)^32,33^ are softer than cardiac tissues (elastic modulus of a few tens of kPa)^34^. Therefore, we downscaled the width and serpentine pitch of the mesh nanoelectronics (ribbon width/thickness = 7.5/1.6 μm) by 25% and 50%, respectively, compared to our previous design^26^ to decrease its effective bending stiffness to 6.7×10^−16^ N.m^2^ (or flexural rigidity of 0.090 N·m), which is more than 50 times lower, and comparable to the mechanical properties of brain tissues^33^. Third, to control the initial cell numbers for brain organoids culture^26^, a polydimethylsiloxane (PDMS) ring (thickness of 100-200 μm, diameter of 6.5 mm) was cast around the mesh nanoelectronics to define the initial region of the stem cell sheet, which ultimately controls the cell number and the size of the 3D organoid. Fourth, brain organoids take months to years to develop and mature; therefore, to enable a stable electrical recording, we used platinum (Pt) black to modify Pt electrodes due to its long-term stability and low impedance compared to other electrode modification methods^35^ (Fig. 1d, Supplementary Fig. 4). The electrochemical impedance of the Pt black-coated sensors (diameter of 25 μm) has an average impedance modulus of (1.40 ± 0.50) × 10^5^ Ω (mean ± S.D., n = 16) at 1 kHz frequency at the day of implantation, and only slightly increased to (3.00 ± 0.33) × 10^5^ Ω after 180 days of incubation in 1x phosphate-buffered saline (PBS). Last, to simultaneously culture multiple brain organoids, each reactor contains four independent 16-channel devices. Electrodes from each device can be individually connected to the voltage amplifier through flip chip-bonded anisotropic conductive film/flat flexible cables (Fig. 1e). Notably, we used photolithography to define a pair of center-symmetric and unique binary fluorescence electronic barcodes (E-barcode) for each sensor by doping 0.005 wt% Rhodamine 6G (R6G) in SU-8 precursors^36^. The fluorescence Ebarcode (Fig. 1d) will be used to determine the 3D position of each sensor within the brain organoids by *post hoc* tissue clearing, staining, and confocal microscopic imaging after electrophysiological recording. We dissociated hiPSC-derived neurons and then cultured them on the stretchable mesh nanoelectronics released on a thin layer of Matrigel. From day 0 to 4 of assembly, bright field (BF) phase images (Fig. 1f) show that hiPSC-derived neurons self-organized into a neuroepithelium-like structure and then gradually folded into a 3D structure with the stretchable mesh nanoelectronics fully embedded. These results proved substantially that the mechanical and structural properties of stretchable mesh nanoelectronics allow for effective embedding and integration into brain organoids through the organogenetic process.

### Long-term electrical recording of human brain organoids

We first built type A cyborg brain organoids to demonstrate the stable long-term electrical recording of human brain organoids. We differentiated hiPSCs on a 2D substrate until they acquired spontaneous electrical activities. Their 2D development and differentiation were assessed by BF phase and fluorescence imaging (Supplementary Fig. 5). After 4 months of differentiation, the 2D hiPSC-derived neurons were dissociated and seeded on a 2D mesh electrode array (of the same design as the stretchable mesh nanoelectronics) for electrophysiological recordings. Spontaneous extracellular action potentials and bursts could be reliably detected and susceptible to glutamate receptor antagonists. Application of 20 μM (2R)-amino-5-phosphonopentanoate (D-AP5) and 20 μM cyanquixaline (CNQX) could significantly reduce the number of spikes and bursts (Supplementary Fig. 6). After confirming the electrical activity, cells were cultured into 3D organoids, which were then placed on a 2D mesh electrode array. Spontaneous extracellular action potentials could be reliably detected from the bottom layer of brain organoids (Supplementary Fig. 7). Both experiments confirmed the electrical activities from the hiPSC-derived neurons generated through our protocol as well as the ability to record single-neuron electrophysiology by the mesh nanoelectronics.

The hiPSC-derived neurons were then dissociated and seeded on the stretchable mesh nanoelectronics/Matrigel hybrids, forming 3D cyborg brain organoids (Fig. 1a, Supplementary Fig. 8). After one-month post-assembly (Fig. 2a), we can detect spontaneous local field potentials (LFPs) and single-unit action potentials from the tissue-wide embedded mesh electrodes (Fig. 2b, Supplementary Fig. 9). To confirm that signals were from neurons, brain organoids were first exposed to the 30 mM potassium chloride (KCl) solution to induce neuronal membrane depolarizations. A statistically significant increase of the electrical activity (1,128 ± 1,894%, mean ± S.D., signal root mean square (RMS) amplitude, n = 12 channels, p<0.01, two-tail, paired t-test) could be recorded from the cyborg brain organoids (Supplementary Fig. 10a, b). Then, the cyborg brain organoids were exposed to glutamate receptor antagonists (20 μM CNQX and 20 μM D-AP5 solution). A statistically significant decrease (50.4 ± 18.6%, mean ± S.D., decrease in signal RMS amplitude, n =12 channels, p<0.01, two-tail, paired t-test) of the electrical activity could be recorded from cyborg brain organoids (Supplementary Fig. 10a, c).

**Figure 2.**
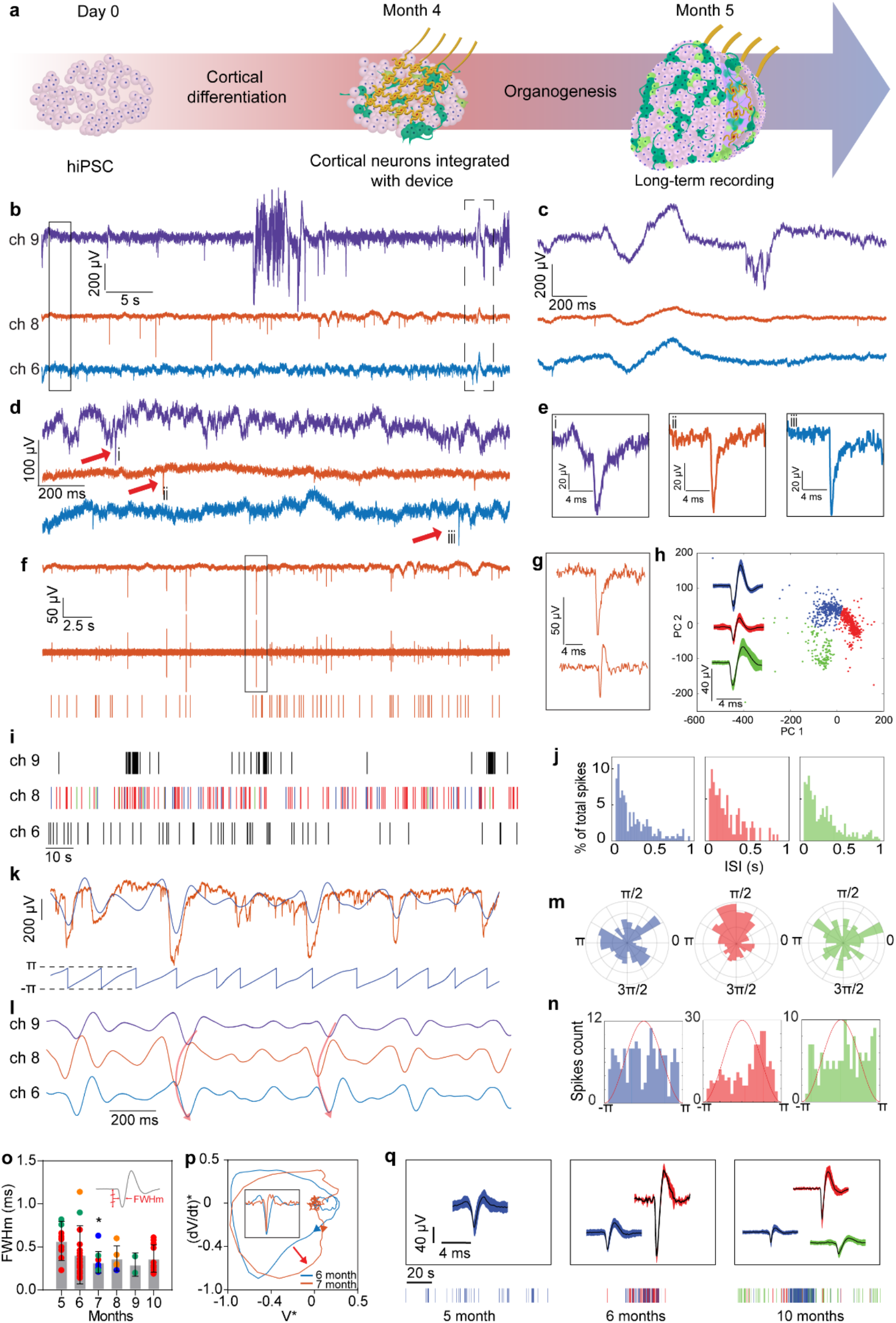
Long-term electrical recording of human brain organoids. **a**, Schematic of the stepwise assembly of mesh nanoelectronics with cells for type A cyborg brain organoids. **b**, Raw voltage traces of 3 representative channels showing neural activities at month 5 of differentiation (*i.e*., month 1 of assembly). **(c, d)** Zoom-in panels from the black box (**c**) and dashed box (**d**) highlighted regions in (**b**) showing that local field potential (LFP) and action potentials recorded are independent between each channel. **e**, Zoom-in window on the single-unit action potentials detected from each channel in (**d**) highlighted by red arrows. **f**, Raw trace (top) and corresponding filtered trace in the frequency band of 300 – 3000 Hz (middle) and raster plot (bottom) from representative channel 8 in (**b**). **g**, Zoom-in window on the same spike before (top) and after filtering (bottom) from the black box highlighted region in (**f**). **h**, Clustering results from the principal component analysis (PCA) applied to the filtered trace in (**f**). Inset shows the average waveforms for each cluster (± S.D.). **i**, Raster plot of the 3 channels represented in (**b**). Colors in the raster plot of channel 8 correspond to the different neurons detected in (**h**). **j**, Interspike intervals (ISI) histogram for each of the neurons in (**h**). **k**, Top: Raw trace (orange) and superimposed theta waves (4-8 Hz) extracted from the same channel (blue). Bottom: Corresponding theta angle. **l**, Theta waves of the channels 6, 8, and 9 shown in (**b**), Red arrows qualitatively show the time delay between each channel. **m**, Circular distribution of theta angles for the firing events of each neuron detected in channel 8. **n**, Distribution of theta angles shown relative to theta cycle (red line). **o**, Full width at half minimum (FWHm) of depolarization of each neuron detected on each cyborg brain organoid. Colors correspond to the data from the same organoid (n=4) over time as a function of culture time (mean ± S.D., *P< 0.05 using one-way ANOVA with the month 5 group as control). **p**, Normalized phase plot of single-unit action potentials (inset) detected on the same channel at months 6 and 7 post-differentiation, showing an increase in the depolarization’s speed. **q**, Single-unit action potentials (top) and corresponding raster plots (bottom) detected from the same channel at 5, 6, and 10 months of culture.

Representative voltage traces showed the temporal delays between LFPs from different channels, suggesting the ability of tissue embedded electrodes to record global signal propagation across the brain organoids (Fig. 2c). Moreover, both burst firing (Supplementary Fig. 11) and individual action potentials (Fig. 2d-e) could be detected from the tissue-embedded electrodes, which were consistent with previous results from 2D MEA^1^, demonstrating the capability to capture the activities from the coordinated neural network and individual neurons. We carried out statistical analysis on the single-unit action potentials to study the evolution of neuronal activities over the time course of brain organoid development. We filtered the signals by 300-3000 Hz bandpass filter (Fig. 2f, g) and applied spike sorting to extract single spikes for analysis. From the representative channel, three single-unit action potentials could be identified through principal component analysis (PCA) (Fig. 2h-j), showing multiple neurons can be detected from each channel. We noticed that the total duration of single-unit action potentials from brain organoids were slower (2- to 3-ms duration) compared with the duration of action potentials detected by mesh nanoelectronics from adult animal’s brain (1- to 2-ms duration)^37,38^, which may suggest the immature nature of neurons in brain organoids. We characterized the propagation of the neural activity across the organoid by analyzing theta waves (4-8 Hz frequency band) (Fig. 2k). A clear temporal delay across channels could be detected (Fig. 2l, Supplementary Fig. 12), which agrees with previous reported LFP detection from brain organoids, further demonstrating the capability of 3D embedded mesh electrodes to map the tissue-wide electrical activity propagation. We did not observe a consistent phase-locking of single-unit action potentials to theta oscillations (Fig. 2m, n)^22,39,40^, which may be due to the immaturity in the connectivity between neurons within the organoids. But the proximity and high density of neurons surrounding the tissue-embedded sensors allowed us to observe the field potentials emerge from collective neuronal activities. We observed a strong correlation between field potential events and spiking bursts (Supplementary Fig. 13).

To confirm the capability of mesh nanoelectronics to chronically record the functional development of the brain organoids at single-cell resolution, we conducted a 6-month electrical recording from the same brain organoids (n = 4 organoids) and performed statistical analysis of single-neuron signals. Long-term recording showed that the average full width at half minimum (FWHm) of the depolarization of action potential decreased from 0.57±0.23 ms in month 5 of differentiation (n = 27 single units, mean ± S.D.) to 0.37±0.16 ms in month 10 of differentiation (n = 11 single units, mean ± S.D.) (Fig. 2o), while the one-way analysis of the variance (ANOVA) suggested that this change is not statistically significant. Similarly, we found that the spiking rates were not significantly changing over time (Supplementary Fig. 14). Using the phase-space^41^ analysis to characterize the evolution of single-unit action potential waveform, the result suggests that the membrane depolarization rate has been increased throughout the development of brain organoids (Fig. 2p). In addition, the waveforms clustering from representative channels shows that the detected number of single-unit action potential increases over development (Fig. 2q). The electrical measurement from the type A cyborg brain organoid system proved the capability to continuously record 3D signals from brain organoids. The results suggested a minimal change of organoid-wide neuronal activities during the 6-month 3D culture in type A brain organoids.

### Tracing of electrophysiology during early brain development

We applied the cyborg organoid platform to early-stage brain organoids, aiming to demonstrate the ability to capture the emergence of single-neuron action potentials. The stretchable mesh nanoelectronics was integrated with hiPSCs before neural differentiation (Fig. 1a) to build type B cyborg brain organoids. To investigate the effects of the embedded mesh nanoelectronics on cell differentiation, type B cyborg brain organoids at different differentiation stages were fixed, sectioned, and immunostained for stage-specific protein marker expressions and compared to control brain organoids without mesh nanoelectronics embedding (Fig. 3a). From day 40 to 90 of differentiation in type B brain organoids, the density of neurons, cortical progenitors, and neural progenitors was consistent across different samples, as shown by the consistent Hexaribonucleotide Binding Protein-3 (NeuN) and beta Tubulin 3 (Tuj1), Paired-box 6 (Pax6), and Nestin expression levels, respectively. Cortical neuron density statistically significantly increased during the course of development, as shown by the increase of cells expressing T-box brain protein 1 (TBR1), which suggests the continuous differentiation and development of neurons over time. Importantly, statistical results showed no significant difference in different types of cells between cyborg and control organoids, confirming the minimal interruptions from the implanted mesh nanoelectronics to the neural differentiation in brain organoids (Fig. 3b).

We started to record the neural activities from type B cyborg brain organoids (n=7 organoids) after one-month differentiation and organogenesis when the 3D organoids were formed (Fig. 4a). A gradually increased activity could be detected from brain organoids over the first 3 months post-differentiation (Fig. 4b). To analyze the single-neuron signals from early developmental stages, we filtered voltage traces in the range of 100-3000 Hz, instead of 300-3000 Hz, which retained slow spikes detected at months 1 and 2 post-differentiation (Supplementary Fig. 15). The results showed that both signal amplitude and the number of single-unit action potentials detected per channel increased over time (Fig. 4b). Moreover, spectral analysis from representative channels showed an increase in power between 0 and 1.5 kHz during the first three months of differentiation (Fig. 4c, d). Statistical analysis of all the cyborg brain organoids showed the significant increase in the power at 300 Hz from month 1 to 3 post-differentiation (n=16 channels, p<0.01, one-way ANOVA) (Fig. 4e), suggesting the gradually increased neuronal activities and functional development of brain organoids, which agrees with the protein marker staining results in Figure 3.

**Figure 3.**
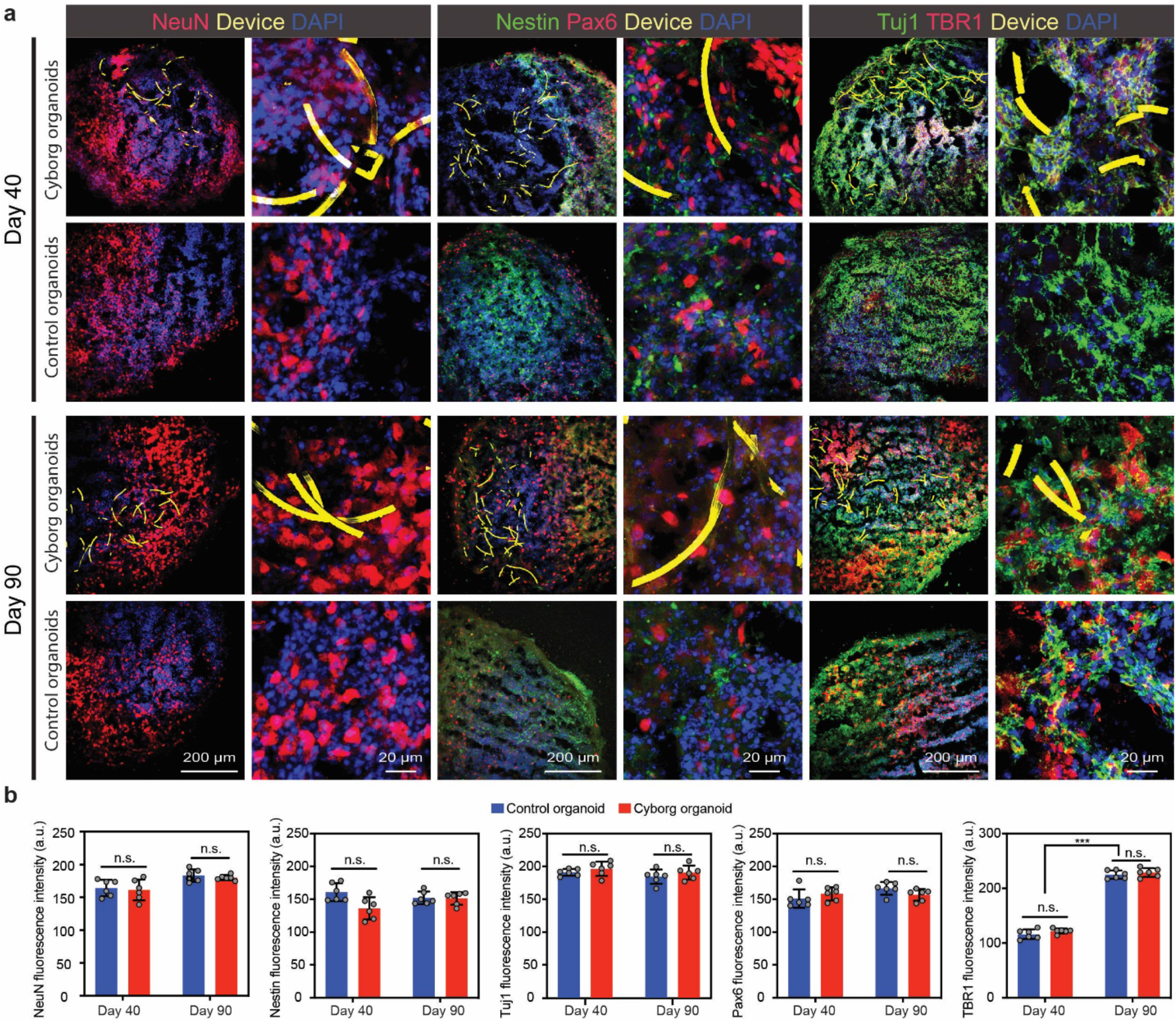
Fluorescence images of cyborg brain organoids. **a**, Immunostaining and fluorescence imaging characterize the development of neurons in type B brain organoids at day 40 and 90 of differentiation with and without the mesh nanoelectronics. **b**, Averaged fluorescence intensity of NeuN, Nestin, Tuj1, Pax6, and TBR1 in the organoid at day 40 and 90 of differentiation with and without the mesh nanoelectronics. (mean ± s.e.m., n=6, *** P< 0.0001, two-tailed, unpaired t-test.)

Next, we analyzed the spike waveform of single-unit action potentials. Phase-space analysis of individual spikes detected from the same electrode showed an increase in depolarization rate from months 2 and 3 (Fig. 4f). Statistical analysis of FWHm and spiking rate of each single-unit spike (n = 7 organoids) showed the narrowing of spikes and increase in spiking rate over time (Fig. 4g, h). FWHm decreased significantly from 1.05±0.63 ms in month 1 (n = 9 single units, mean ± S.D.) to 0.38±0.22 ms in month 3 (n = 15 single units, mean ± S.D.). Plotting spiking rate per neuron as a function of FWHm^−1^ further revealed the positive correlation between narrowing and spiking activity of single units. While the increase in mean firing rate and signal power has been previously reported^1,21,25^, the narrowing of spikes’ depolarization in brain organoids generated from hiPSCs is reported for the first time due to the ability to form direct contact between electrodes and neurons in 3D brain organoids. This result also agrees with previous findings for hiPSC-induced 2D neuron cultures^42^.

**Figure 4.**
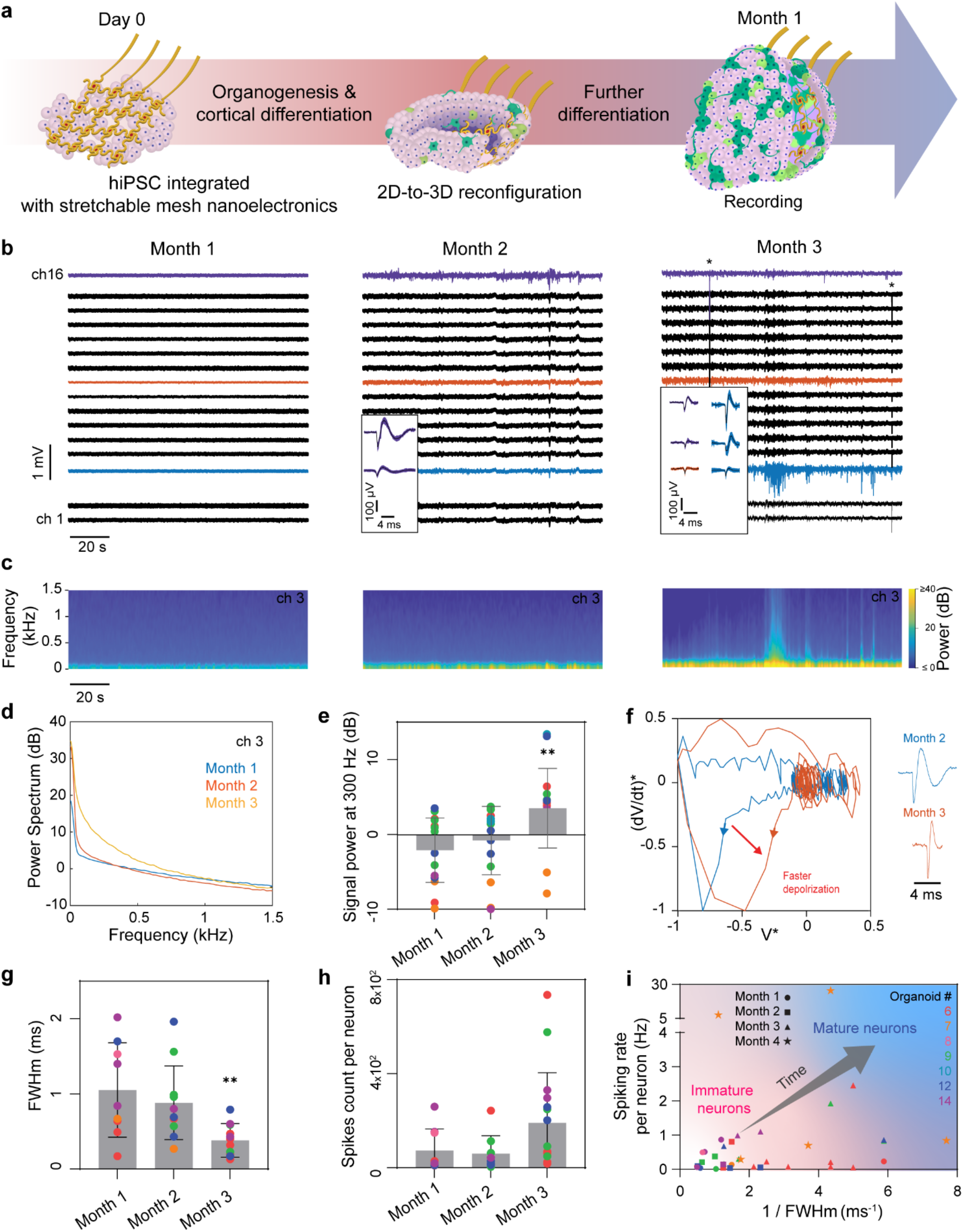
Electrical recording of human brain organoids during early development. **a**, Schematics of the stepwise assembly of mesh nanoelectronics with hiPSCs for type B cyborg brain organoids. **b**, Raw voltage traces at 1, 2, and 3 months after cortical differentiation. Single-spike waveforms extracted from the voltage traces filtered by 100 Hz high-pass filter are plotted as insets. Colors match with the corresponding voltage trace of the channel. **c**, Spectrograms at 1, 2, and 3 months of differentiation for channel 3 showing a strong increase in power between 0 to 1 kHz after 3 months of differentiation. **d**, Corresponding power spectrums to (**c**). **e**, Signal power at 300 Hz for electrodes with detected neural activity. Colors correspond to the data from the same organoid (n=7) over time. **f**, Normalized phase plot of single-unit action potentials and its corresponding waveforms (inset) detected from the same channel at 2 and 3 months of differentiation, showing an increase in the rate of depolarization. **g-h**, FWHm of depolarization (**g**) and spikes count per neurons per 2-min recording (**h**) at 1, 2, and 3 months of differentiation. Colors correspond to the data from the same organoid (n=7) over time. **i**, Spiking rate per neuron detected as a function of the inverse of the FWHm of depolarization, showing that the single-cell action potential spikes from neurons evolve towards shorter spike width and higher spiking rate over the time course of brain organoid development. In panels (**e, g**), value=mean ± S.D., ** P<0.01, one-way ANOVA with “Month 1” group as control.

To confirm that spiking activity arises from synaptic transmissions and diverse types of neurons throughout development, we applied GABAergic (bicuculline) and glutamatergic (CNQX and D-AP5) receptor antagonists to block inhibitory and excitatory synaptic transmission. Results showed that application of 10 μM bicuculline (BCC) can introduce a statistically significant increase of 2,870 ± 2,575% (mean ± S.D.) in spontaneous spiking rate (n=4 organoids, p<0.05, two-tailed, paired t-test), while application of 20 μM CNQX and 20 μM D-AP5 can introduce a statistically significant decrease of 91 ± 16% (mean ± S.D.) in spontaneous spiking rate (n = 4 organoids, p<0.05, two-tailed, paired t-test) (Supplementary Fig. 16). These results suggest that both inhibitory and excitatory connections have been built in the brain organoids after 3-month neuronal differentiation.

We applied the intact organoid clearing and imaging to confirm the seamless integration of the stretchable mesh electrodes (Supplementary Fig. 17) with type B cyborg brain organoids. Each electrode channel in 3D could be identified through imaging and identifying the fluorescent E-barcodes paired with electrodes, and imaged with the density of neurons around the electrodes where strong neural activities were detected (Fig. 5a). Indeed, a high density of neurons could be observed at the vicinity of such sensors (*i.e*. channels 3 and 9) (Fig. 5b, Supplementary Fig. 18). As in type A brain organoids, we could also capture LFP, non-correlated individual action potentials, and spontaneous bursting activity from type B cyborg brain organoids at 3 months post-differentiation (Fig. 5c-e, Supplementary Fig. 19), which agrees with the results from the previous report^1^. Spike sorting revealed that multiple neurons can be measured by the same channel (Fig 5.f-j), and theta-wave analysis showed a clear temporal delay across channels, although phaselocking of neurons firing was not observed (Fig. 5.k-n)^22^. Finally, statistical analysis showed that while the FWHm of spikes’ depolarization of type B cyborg brain organoids at 1 month postdifferentiation (1.05±0.63 ms, n=9 single units, mean ± S.D.) is significantly longer than other samples, the FWHm at 3 months post-differentiation (0.38±0.22 ms, n = 15 single units, mean ± S.D.) is not statistically different to the FWHm of type A cyborgs at 5 months post-differentiation (0.57±0.23 ms, n = 27 single units, mean ± S.D. two-tailed, unpaired, t-test) (Fig. 5o). These results suggest that the cyborg brain organoid platform can be used to quantify the electrophysiological evolution of neurons during the development phase and detect variations across samples prepared by different protocols.

**Figure 5.**
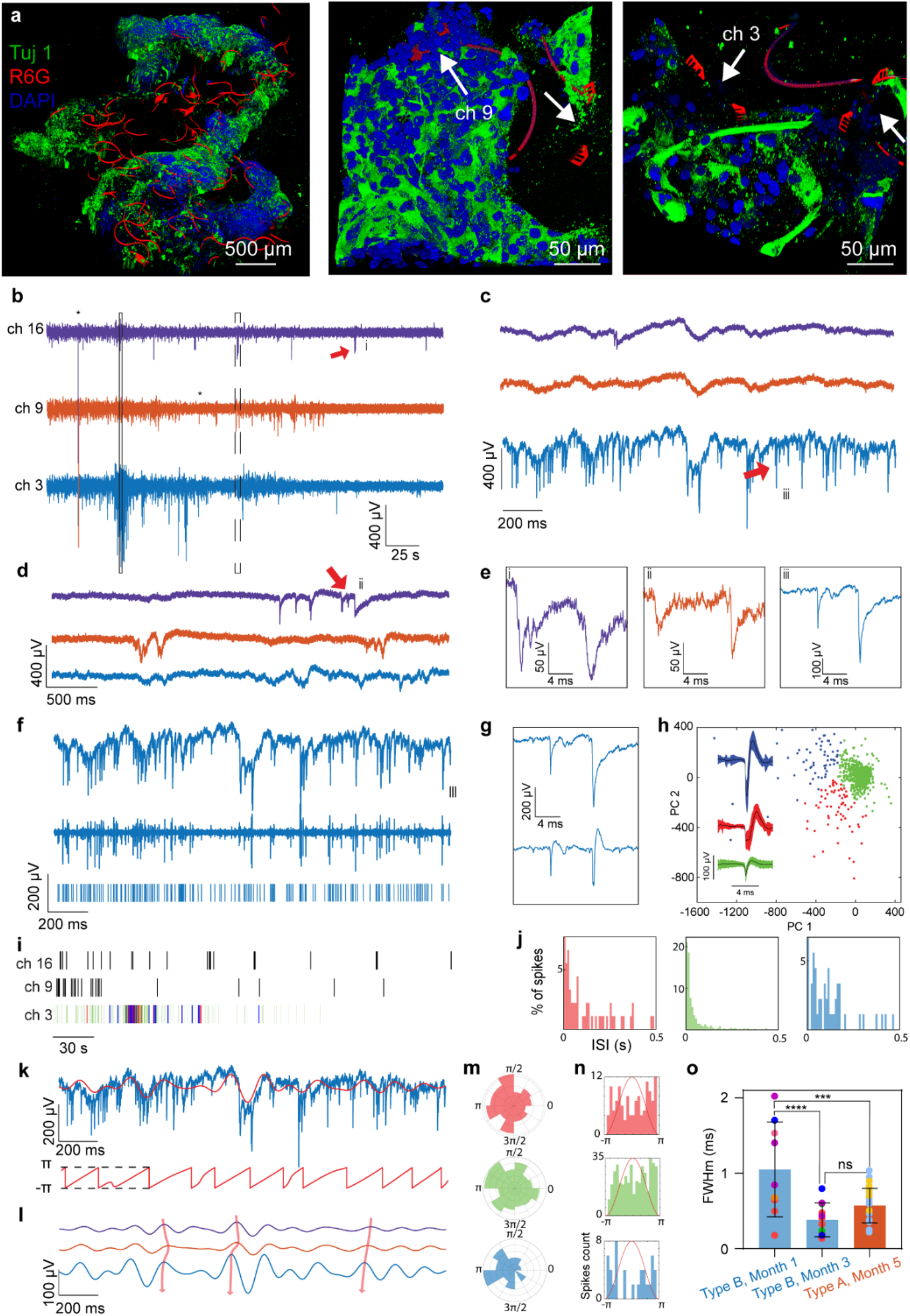
Electrophysiology of human brain organoids during early developmental stage. **a**, 3D views of reconstructed fluorescence images of cleared, immunostained type B cyborg brain organoids at month 3 of differentiation. Red, green, and blue colors correspond to R6G, Tuj 1, and 4’,6-diamidino-2-phenylindole (DAPI). White arrows highlight the position of the sensors. Channel number was read out through the fluorescence electronic barcode identification. **b**, Raw voltage traces of 3 representative channels showing the emergence of neural activities at 3 months post-differentiation. * denotes voltage artifact (instantaneous large voltage variations synchronous on all channels). **(c, d)**, Zoom-in panels from the black box (**c**) and dashed box (**d**) highlighted regions in (**b**) showing that field potential and action potentials recorded are independent between each channel. **e**, Zoom-in views of single-unit action potentials detected from (**b-d**) highlighted by red arrows. **f**, Raw trace (top) and corresponding filtered trace in the frequency band of 100-3000 Hz (middle) and corresponding raster plot (bottom). **g**, Zoom-in window on the same spike before (top) and after filtering (bottom). **h**, Clustering results from the PCA applied to the filtered trace in (**f**). Inset shows the average waveforms for each cluster (± S.D.). **i**, Raster plot of the 3 channels represented in (**b**). Colors in the raster plot of channel 3 correspond to the different neurons detected by clustering in (**h**). **j**, ISI histograms for each of the clusters presented in (**h**). **k**, Top: Raw trace (blue) and superimposed theta waves (4-8 Hz) of the same channel (red). Bottom: Corresponding theta angle. **l**, Theta waves of the channels 3, 9, and 16 shown in (**b**), Red arrows show the time delay between each channel. **m**, Circular distribution of theta angles for the firing events of each neuron detected in channel 3. **n**, Distribution of theta angles shown relative to theta cycle (red line). **o**, Comparison of the full width at half minimum (FWHm) of neuron’s depolarization at months 1 and 3 of early-stage (type B brain organoids, n=4) and month 5 (type A brain organoids, n=2) of long-term electrophysiological recordings. Colors correspond to the data from the same organoid. (mean ± S.D., ** P<0.01 and **** P<0.0001, one-way ANOVA with the group “Type B, month 1” as control).

## Discussion

We have demonstrated a cyborg brain organoid platform that can stably record 3D brain organoid-wide, millisecond-timescale, and single-neuron electrophysiology over the time course of brain organoid development. Long-term stable electrophysiological measurements of neural activity throughout the development of brain organoids reveal the increased number of neurons that can be detected from electrodes and millisecond-timescale evolution of the single-unit action potential waveform over 3D development. In addition, electrophysiological measurements during early brain organoid development reveal that single-unit action potentials can be detected from brain organoids after 3-month neuronal differentiation. The increase of action potential amplitude, synchrony, and firing rate as well as narrowing of action potential duration suggest changes in neural network connectivity and cellular ion channel expression level. We envision that further integration of cyborg brain organoids with organoid-wide connectomics and spatial transcriptomics mapping will further test this hypothesis^36,43^.

Cyborg brain organoid technology can potentially become a useful tool for quantifying the functional development of brain organoids and standardizing the culture conditions across different types of protocols by tracing organoid-wide tissue and single-cell electrical activity during the entire organoid development. It will also be useful for studies of developmental neuroscience and drug screening. As the stretchable mesh nanoelectronics are fabricated by the standard lithographic process, this method is scalable by integrating a larger number of sensors and stimulators through the integration of multiplexing integrated circuits (IC)^44–46^. Further integration of multifunctional sensors and stimulators will offer the potential of combined multimodal interrogation (*e.g*., mechanical and chemical) and intervention (*e.g*., optogenetics) capabilities to the cyborg brain organoids platform for human brain developmental studies.

## Online methods

### Device fabrication

Fabrication of the ultra-thin, stretchable mesh nanoelectronics made of SU-8 negative photoresist was based on methods described previously^18^. The key steps (Supplementary Fig. 1) included: (**1**) Cleaning a glass wafer (500-nm thickness) with acetone, isopropyl alcohol, and water. (**2**) Depositing 100-nm-thick nickel (Ni) using a thermal evaporator (Sharon) as a sacrificial layer. (**3**) Spin-coating SU-8 precursor (SU-8 2000.5, MicroChem, 800-nm or 400-nm thickness for, respectively, a spinning at 1000 or 3000 rpm), followed by pre-baked at (65 °C, 95 °C) for 2 min each, exposed to 365 nm ultra-violet (UV) for 200 mJ/cm^2^, post-baked at (65 °C, 95 °C) for 2 min each, developed using SU-8 developer (MicroChem) for 60 s, and baked at 180 °C for 40 min to define mesh SU-8 patterns (800-nm or 400-nm thickness for, respectively, spinning at 1,000 or 3,000 rpm) for bottom encapsulation. To define the fluorescent electronic barcode, 0.005 wt% of Rhodamine 6G powder (Sigma-Aldrich) was added into the SU-8 precursor. (**4**) Spin-coating LOR3A photoresist (MicroChem) at 4,000 rpm, followed by pre-baking at 180 °C for 5 min; spin-coating S1805 photoresist (MicroChem) at 4000 rpm, followed by pre-backing at 115 °C for 1 min; exposed to 405 nm UV for 40 mJ/cm^2^, and developed using CD-26 developer (Micropost) for 70 s to define interconnects patterns. (**5**) Depositing 5/40/5-nm-thick chromium/gold/chromium (Cr/Au/Cr) by electron-beam evaporator (Denton), followed by a standard lift-off procedure in remover PG (MicroChem) overnight to define the Au interconnects. (**6**) Repeating Step (**4**) to define sensors tip patterns in LOR3A/S1805 bilayer photoresists; (**7**) Depositing 5/50-nm-thick chromium/platinum (Cr/Pt) by electron-beam evaporator (Denton), followed by a standard lift-off procedure in remover PG (MicroChem) overnight to define the Pt electrodes. (**8**) Repeating Step (**3**) for top SU-8 encapsulation and SU-8 barcodes (containing R6G at a concentration of 50 μg/mL). (**9**) Electroplating Pt black or polymerizing PEDOT on the Pt electrode using the recipe described below. (**10**) Soldering a 16-channel flexible flat cable (Molex) onto the input/output pads using a flip-chip bonder (Finetech Fineplacer). (**11**) A 500 μm-thick PDMS ring was attached around each device after 2-min oxygen plasma (Anatech 106 oxygen plasma barrel asher) treatment at 50W. (**12**) Gluing a chamber onto the substrate wafer to completely enclose a 2×2 mesh device array using a bio-compatible adhesive (Kwik-Sil, WPI). (**13**) Treating the surface of the device with light oxygen plasma (Anatech 106 oxygen plasma barrel asher), followed by adding 3 mL of Ni etchant (type TFB, Transene) into the chamber for 2 to 4 hours to completely release the mesh electronics from the substrate wafer. The device was then ready for subsequent sterilization steps before cell culture.

### Materials preparation

We use an SP-150 potentiostat from Bio-logic© alongside its commercial software EC-lab in voltage or current control for electrodeposition. Electrodes from devices are connected to the working electrode. The counter electrode is a platinum wire, also serving as a voltage reference, which is immersed in the precursor solution. For Pt black electrodeposition, the precursor (0.8 wt% Chloroplatinic acid solution) solution was applied by the current at 1 mA/cm^2^ for 5-10 minutes.

### Device impedance characterization

A three-electrodes setup was used to measure the electrochemical impedance spectrum of the electrodes from each device. Platinum wire (300 μm in diameter, 1.5 cm in length immersed) and a standard silver/silver chloride electrode were used as counter electrode and reference electrode, respectively. The device was immersed in a 1xPBS solution during measurement. The SP-150 potentiostat (Bio-logic) along with its commercial software EC-lab was used to perform the measurements. For each measurement, at least three frequency sweeps were measured from 1 MHz down to 1Hz to obtain statistical results. A sinusoidal voltage of 100 mV peak-to-peak was applied. For each data point, the response to 10 consecutive sinusoids (spaced out by 10% of the period duration) was accumulated and averaged. Optical images of the measurement setup can be found in our previous work.^26^

### Electrophysiological recordings

A Blackrock CerePlex Direct voltage amplifier along with a 32 channels Blackrock μ digital headstage connected to the device was used to record electrical activity from organoids. The headstage-to-device connector (16 channels) was homemade. The organoid culture medium was grounded to the earth and a reference electrode was also inserted in the media, far from the device (distance above 1 cm). Platinum wires were used as ground and reference electrodes. A sampling rate of 30000 samples per second was used. Band-pass filters (Butterworth, 4^th^ order) were applied depending on the analysis performed. MATLAB codes provided by Blackrock were used to convert raw data files into an accessible format. Data were then transferred to Graphpad Prism for post-processing.

### MATLAB analysis

**Spike detection and clustering**. We developed our own MATLAB code to analyze neural signals (code is available at https://github.com/CyBrainOrg). The threshold for spikes detection was set a −5*standard deviation of the filtered (300 – 6000 Hz or 100 – 6000 Hz bandpass) time series and principal component analysis was used for dimension reduction. MATLAB’s “*kmeans*” function was used to cluster the extracted waveforms and exclude noise artifacts. **Phase-space analysis**. Discrete derivatives of average waveforms are calculated. To exclude amplitude variations from the analysis, the average waveform and derivative of the average waveform were normalized by their minimum value. **Spike phase locking to theta waves**. Theta band oscillations are defined here as voltage traces filtered in the band 4 – 8 Hz. The time series theta phase was determined by taking the standard Hilbert transform of the theta band oscillations and calculating the angle between its real and imaginary components^47^. The distribution of theta phases at spiking events was determined for all clusters with at least 25 action potential spikes. We used the Rayleigh criterion to test the non-uniformity of theta phase distributions, using the open source CircStat toolbox^39,40^. We considered that phase-locking to theta waves happens for P-values below 0.05.^22^

### Organoid culture

HiPSC line hiPSCs-(IMR90)-1 were obtained from WiCell Research Institute (Madison, WI, USA) and cultured in a 6-well plate coated with Matrigel (Corning) in Essential 8 medium (life technologies). Authentication and test for the free of mycoplasma were performed by WiCell Research Institute. Neuron differentiation was based on methods described previously^7^ with minor modification. Briefly, hiPSCs at 2D or 3D hiPSC cyborg organoids were induced for cortical neuron differentiation for 11 days in the induction medium containing DMEM/F12 (50%) and Neurobasal (50%) medium. From Day0 to Day11, 1% N_2_ (Life Technologies), 2% B27 (Life Technologies), SB431542 (10 μM, Selleckchem), and LDN193189 (100 nM, Selleckchem) were applied. The cells were maintained in Neurobasal medium which including BDNF (20 ng mL^−1^, PeproTech), GDNF (10 ng mL^−1^, PeproTech), L-ascorbic acid (200 *μ*M, Sigma), dibutyryl-cAMP (0.5 mM, Santa Cruz Biotechnology), and 2.5 μM DAPT (Selleckchem) after Day 11 of differentiation.

### Cyborg organoids integration^26^

Briefly, the released mesh nanoelectronics was rinsed with DI water and then incubator in 70% ethanol at room temperature for 15 minutes to sterilize. The device was sequentially coated with Poly-D-lysine hydrobromide (Sigma) and Matrigel solution (Corning). Finally, about 20 μL liquid Matrigel (10 mg/mL) was added to each well in the cell culture chamber from the device-free side on ice. The cell culture chamber was incubated at 37 °C for at least 30 minutes to cure the Matrigel layer. hiPSCs or hiPSC-derived neurons (1×10^6^ cells) were suspended in a mixture of E8 or neural maturation medium, and then transferred onto the cured Matrigel in each well of the cell culture chamber and cultured at 37 °C, 5% CO_2_.

### Drug test

The brain organoids were recorded in the neural maturation medium. For drug test^1^, 30 mM KCl, 20 μM CNQX, 20 μM D-AP5, 10 μM bicuculline, and 1 μM TTX were applied. In the measurement, baseline recording was performed before and after the addition of chemicals. The sample was rinsed three times with DPBS every time after the drug test before adding fresh medium.

### Organoids imaging

For whole organoids imaging, procedures were adapted from tissue clearing techniques CLARITY^48^ and passive clarity technique (PACT)^49^, the organoids were fixed with 4% PFA at 4 °C overnight and incubated with hydrogel solution (0.25% (w/v) VA-044 and 4% (w/v) acrylamide in PBS) at 4 °C for 24 hours. The samples were placed in an X-CLARITY hydrogel polymerization device for 3-4 hours at 37 °C with −90 kPa vacuum, followed by a wash in PBS overnight before electrophoretic lipid extraction for 24 hours in the X-CLARITY electrophoretic tissue clearing (ETC) chamber. Then immunostaining was performed by staining the primary antibodies (Tuj or TBR1) for 3-5 days and the secondary antibodies for 2-4 days, respectively. The samples were submerged in optical clearing solution overnight and embedded in 2% agarose gel before imaging using Leica TCS SP8 confocal microscopy. For characterization of cyborg brain organoids, the organoids at day 40 and month 3 were fixed with 4% PFA at 4 °C overnight and immersed in 30% sucrose for at least 12 hours. Then the samples were embedded in optimal cutting temperature (OCT) compound and cryostat section of 30-μm-thick slices. Brain organoids without device integration were used as control. For staining, the first antibodies (NeuN, Tuj, Pax6, Nestin, and TBR1) were incubated at 4 °C overnight and the secondary antibodies were stained at RT for 3-4 hours. For 2D, the cells were fixed with 4% paraformaldehyde (PFA) at room temperature (RT) for 15 minutes. Cells were incubated with primary antibodies at 4°C overnight and the secondary antibodies were stained at RT for 1-2 hours. Finally, 4’,6-diamidino phenylindole 8 (DAPI) were stained for 10 minutes. All samples were imaged by Leica TCS SP8 confocal microscopy. Imaging was analyzed by Leica Application Suite X (LAS X) and Fiji. Fluorescence intensity was calculated by Fiji. Data analysis and statistical tests were performed by Graphpad Prism.

## Data availability

The authors declare that all data supporting the findings of this study are available within the paper and its Supplementary Information.

## Acknowledgements

We acknowledge the discussion and assistance from all Liu Group members. We acknowledge the support from the NIH/NIMH 1RF1MH123948; NSF through the Harvard University Materials Research Science and Engineering Center Grant #DMR-2011754; Harvard University Center for Nanoscale Systems supported by the NSF; Aramont Fund for Emerging Science Research; and the William F. Milton Fund.

## Author contributions

P.L.F., Q.L. and J.L. designed experiments. P.L.F. and R.L. designed, fabricated, and characterized the properties of the stretchable mesh nanoelectronics. Q.L. and H.J. performed cell cultures and device implantations. Q.L., Z.L. and H.J. performed immunofluorescence staining and analyzed related data. P.L.F., Q.L. and S.Z. performed electrophysiological recordings. P.L.F. and Q.L. performed drug tests. P.L.F. and K.T. analyzed electrophysiological recordings. P.L.F., Q.L., K.T. and J.L. wrote the manuscript. J.L. supervised the study.

