## Supplementary figures for "A method for three-dimensional single-cell chronic electrophysiology from developing brain organoids"

### Table of contents

**Figure S1** | Fabrication flow of the stretchable mesh nanoelectronics.

**Figure S2** | Mesh design for nanofabrication.

**Figure S3** | Optical bright-field (BF) imaging of mesh nanoelectronics.

**Figure S4** | Electrochemical impedance characterizations.

**Figure S5** | Fluorescence imaging of 2D human induced pluripotent stem cell (hiPSC) derived neurons culture for the preparation of type A cyborgs.

**Figure S6** | Drug response for 2D neurons culture on non-released microelectrode array.

**Figure S7** | Localization of neural signals under brain organoid cultured on a non-released microelectrode array.

**Figure S8** | Imaging of type A cyborg brain organoids.

**Figure S9** | Raw voltage recording of type A cyborg brain organoid.

**Figure S10** | Type A cyborg brain organoids drug response.

**Figure S11** | Spontaneous bursting activity in type A cyborg brain organoids.

**Figure S12** | Theta oscillations (4 – 8 Hz bandpass) across all channels of the same device.

**Figure S13** | LFP and spiking burst co-activation in a type B cyborg brain organoid.

**Figure S14** | Spiking rate of neurons detected in type A cyborg brain organoids.

**Figure S15** | Effect of 100 – 6000 Hz and 300 – 6000 Hz bandpass filters on spike detection.

**Figure S16** | Type B cyborg brain organoids drug response.

**Figure S17** | 3D reconstructed fluorescence images of a cleared, immunostained type B cyborg brain organoid at month 3 of differentiation.

**Figure S18** | Raw voltage recording of type B cyborg brain organoid.

**Figure S19** | Spontaneous bursting activity in type B cyborg brain organoids.

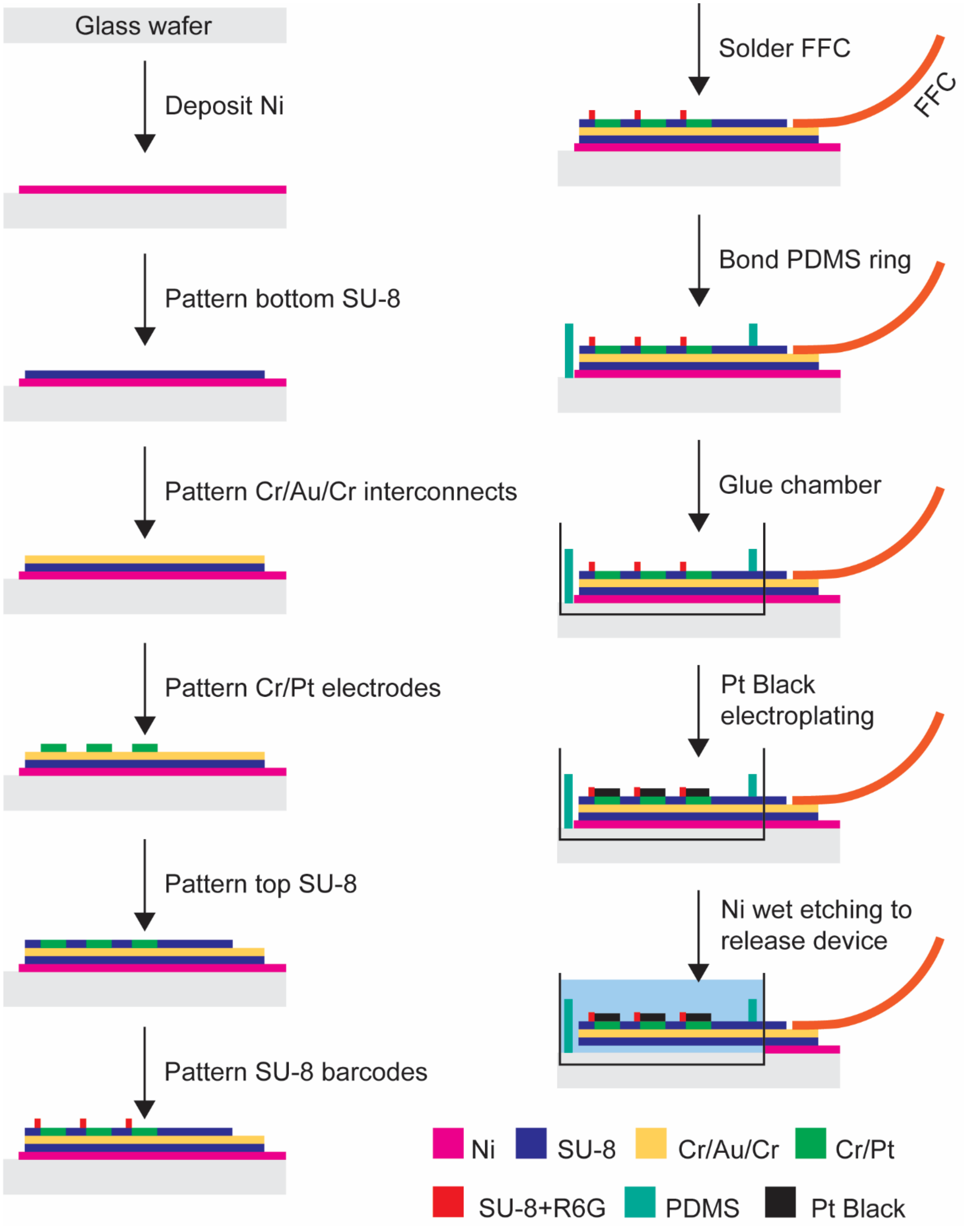


**Figure S1. Fabrication flow of the stretchable mesh nanoelectronics.**

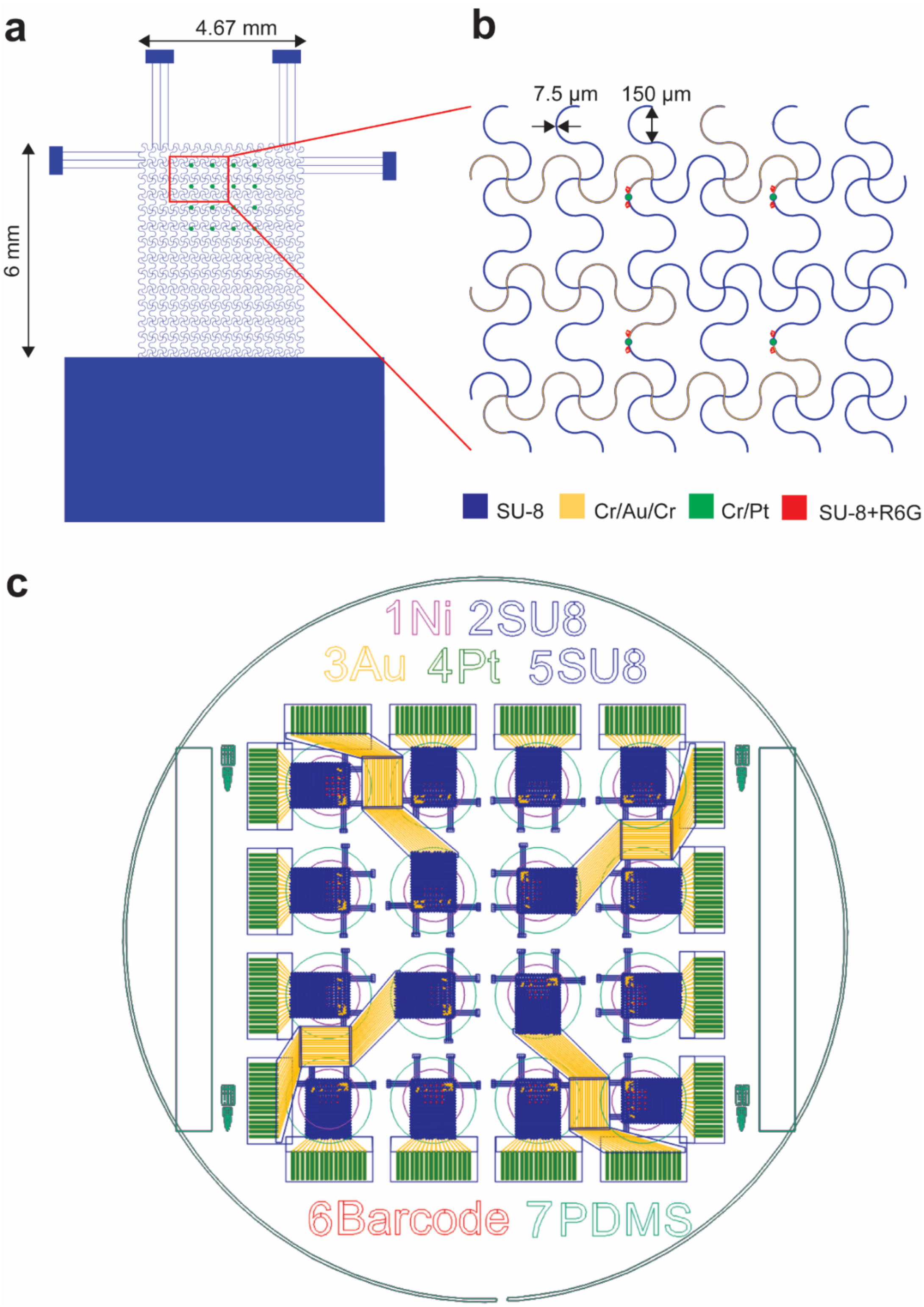


**Figure S2. Mesh design for nanofabrication. a,** Bottom SU-8 encapsulation of a single mesh device. Platinum electrodes (green) are scaled up for visualization. **b**, Zoom-in on an area containing 4 sensors. **c**, 16 devices with 16 electrodes each can be fabricated on a single 3 inch-wafer. The wafer can be diced in 4 quadrants containing 4 devices each.

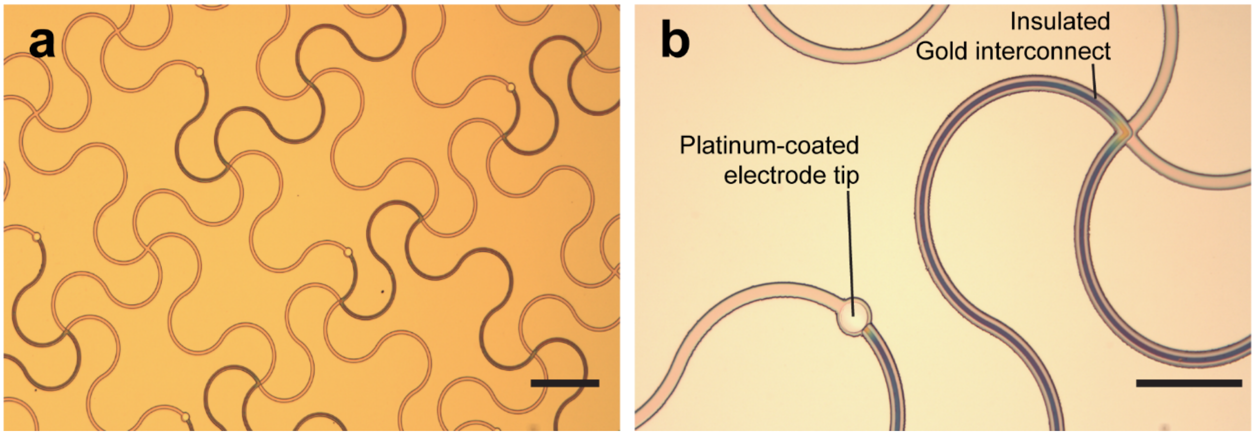


**Figure S3. Optical bright-field (BF) imaging of mesh nanoelectronics.** Images were taken after Platinum deposition on electrode tips and before patterning the SU-8 barcodes. Scale bars 125 µm (**a**), 100 µm (**b**).

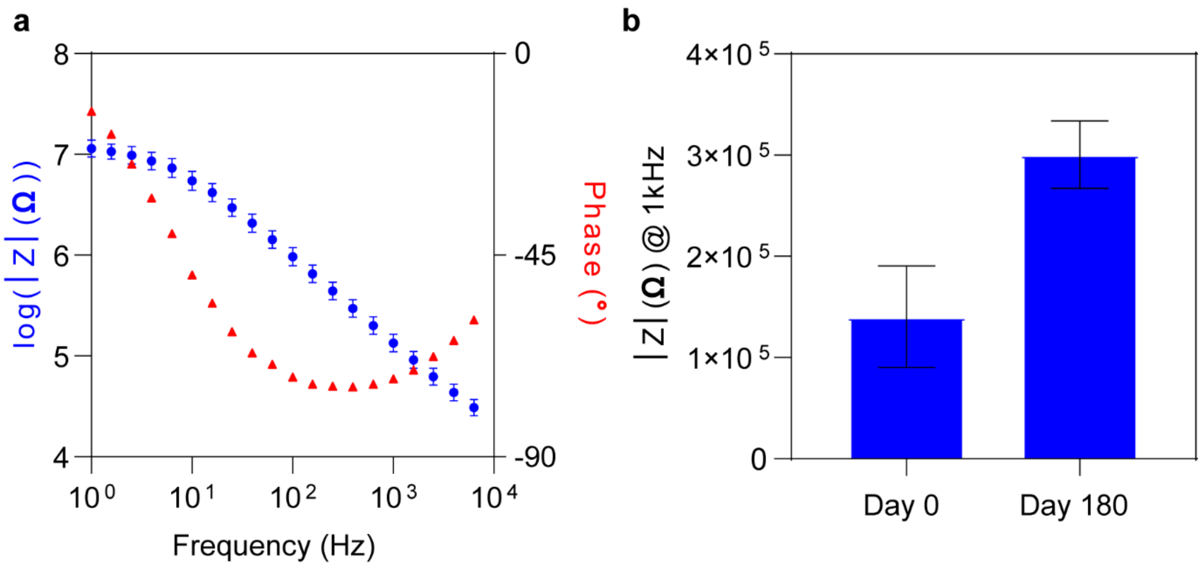


**Figure S4. Electrochemical impedance characterizations. a**, Average Bode plots for electrodes electroplated with Platinum Black (mean ± S.D., n = 4). **b**, Electrochemical impedance modulus of electrodes at 1 kHz. The impedance of Platinum black-coated electrodes increases after 180 days of implantation but remains acceptable for neural signal recordings (mean ± S.D., n = 16).

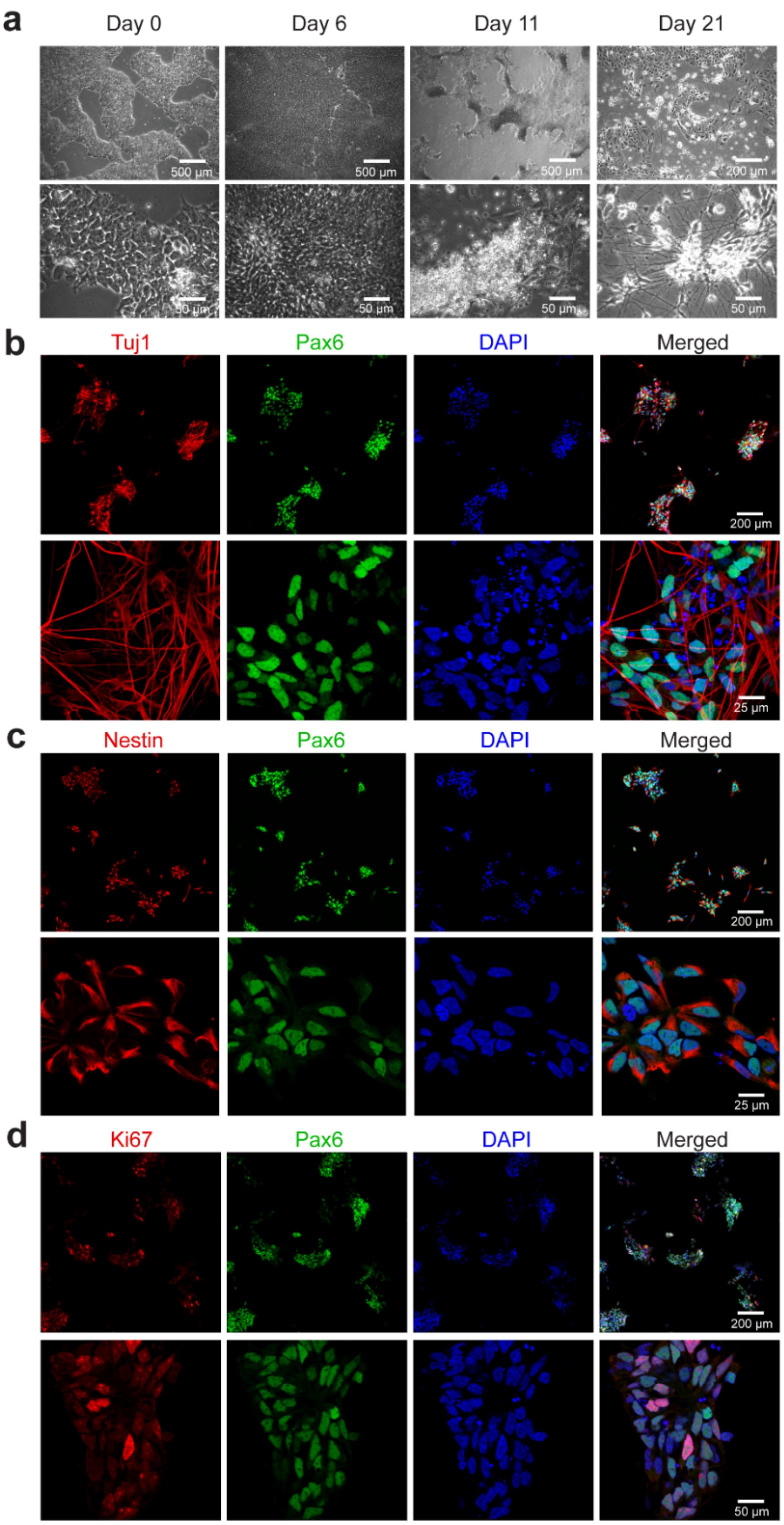


**Figure S5. Fluorescence imaging of 2D human induced pluripotent stem cell (hiPSC) derived neurons culture for the preparation of type A cyborgs. a**, Bright-field (BF) imaging of cells at day 0, 6, 11 and 21 of differentiation. **b-d**, Fluorescence imaging of immunostained 2D neurons at day 20 of differentiation.

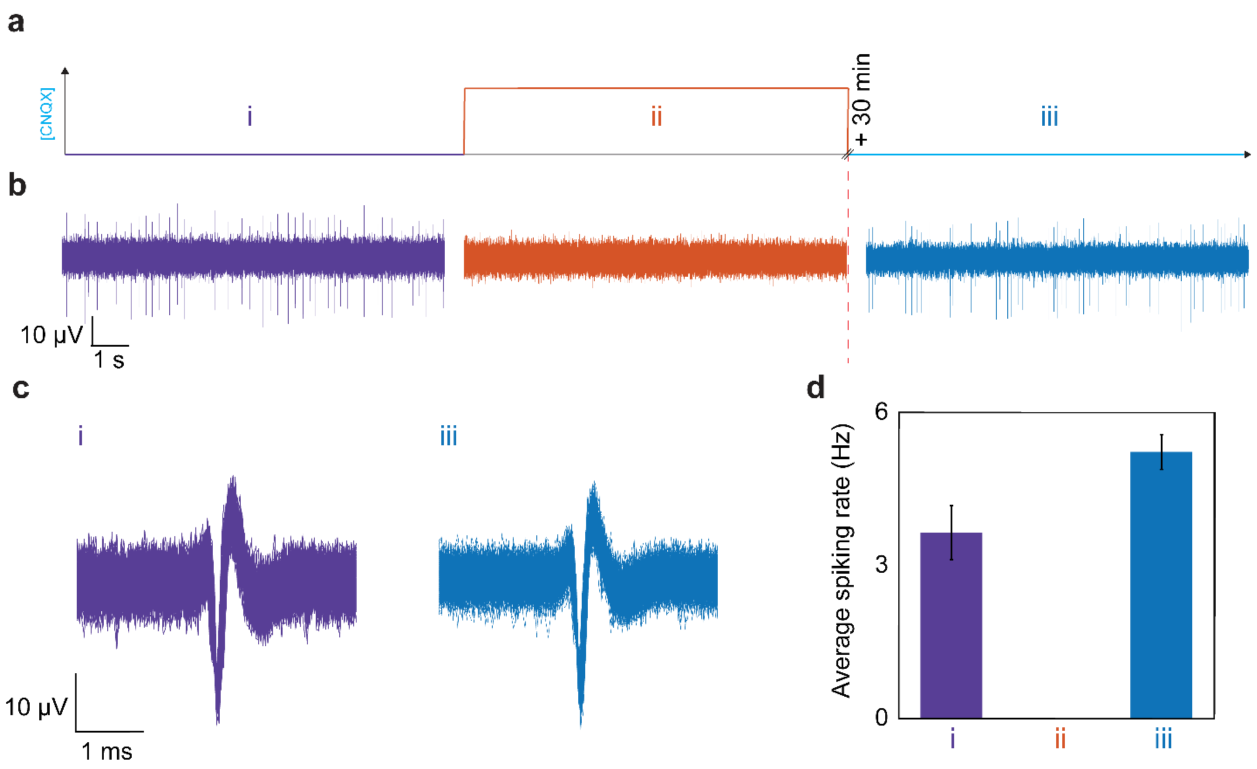


**Figure S6. Drug response for 2D neurons culture on non-released microelectrode array. a**, Schematic shows the protocol of CNQX/D-AP5 injection and washing. The baseline activity of neuron activities is first recorded (i), followed by an injection of CNQX/D-AP5 at 20 µM concentration (ii), and after the culture medium is washed neuron activity is recorded again after 30-min recovery (iii). **b**, Representative voltage trace (filtered in the range of 300-3000 Hz) in step (i), (ii) and (iii). **c**, Waveforms detected in (i) and (iii) show the same features, indicating the recording from the same neuron before and after CNQX/D-AP5 application. **d**, Average spiking rate (mean ± S.D. of the moving average with time bin of 10s) during the three states, showing that CNQX/D-AP5 completely inhibits the activity of the 2D neurons.

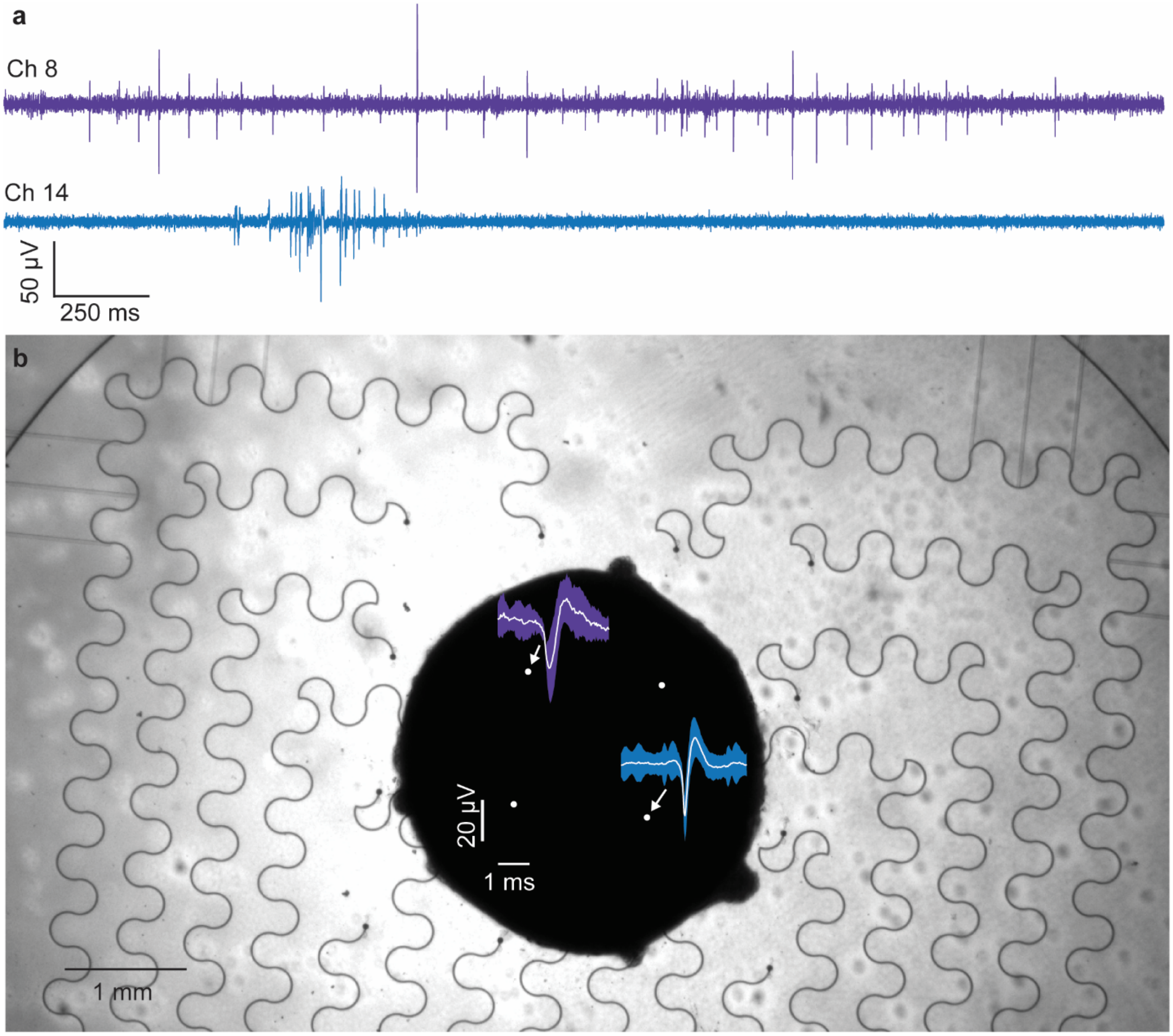


**Figure S7. Localization of neural signals under brain organoid cultured on a non-released microelectrode array. a**, Filtered traces (300-3000 Hz bandpass) of representative electrodes with neural activity. **b,** Waveforms (mean ± S.D.) from voltage recordings in (**a**) are superimposed to the phase imaging, showing that the detected neural signals are measured underneath the brain organoid, where its distance to the non-released mesh device is minimal. Other electrodes did not capture neural signals. The brain organoids were culture in suspension for about 3 month and plated onto the 2D device.

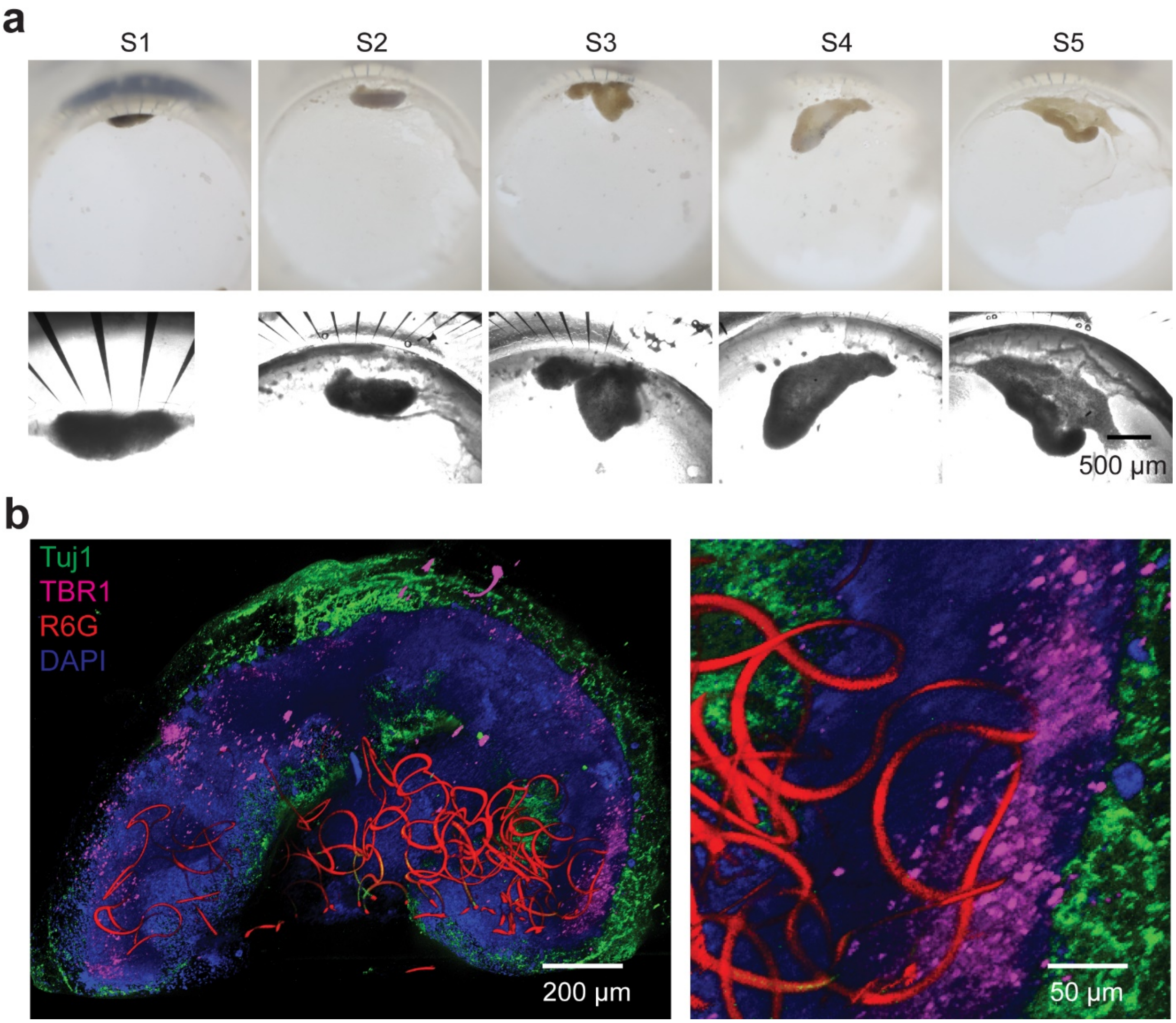


**Figure S8**. **Imaging of type A cyborg brain organoids.** **a**, BF and phase imaging of representative type A cyborg brain organoids one-month post-assembly. **b**, 3D reconstructed fluorescence images of a cross-section of cleared, immunostained type A cyborg brain organoid.

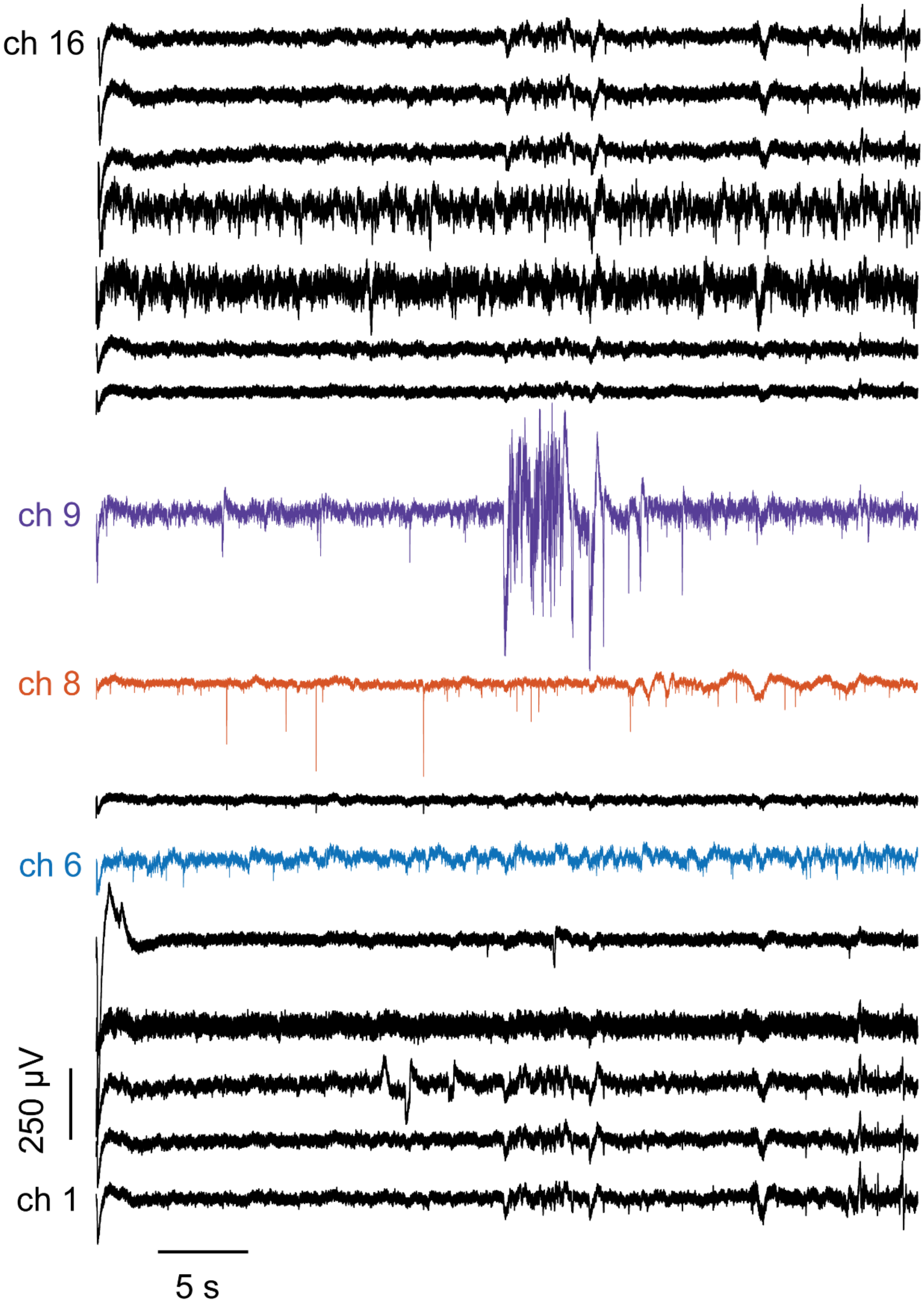


**Figure S9. Raw voltage recording of type A cyborg brain organoid.** Channel 6, 8 and 9 traces are plotted in Figure 2.

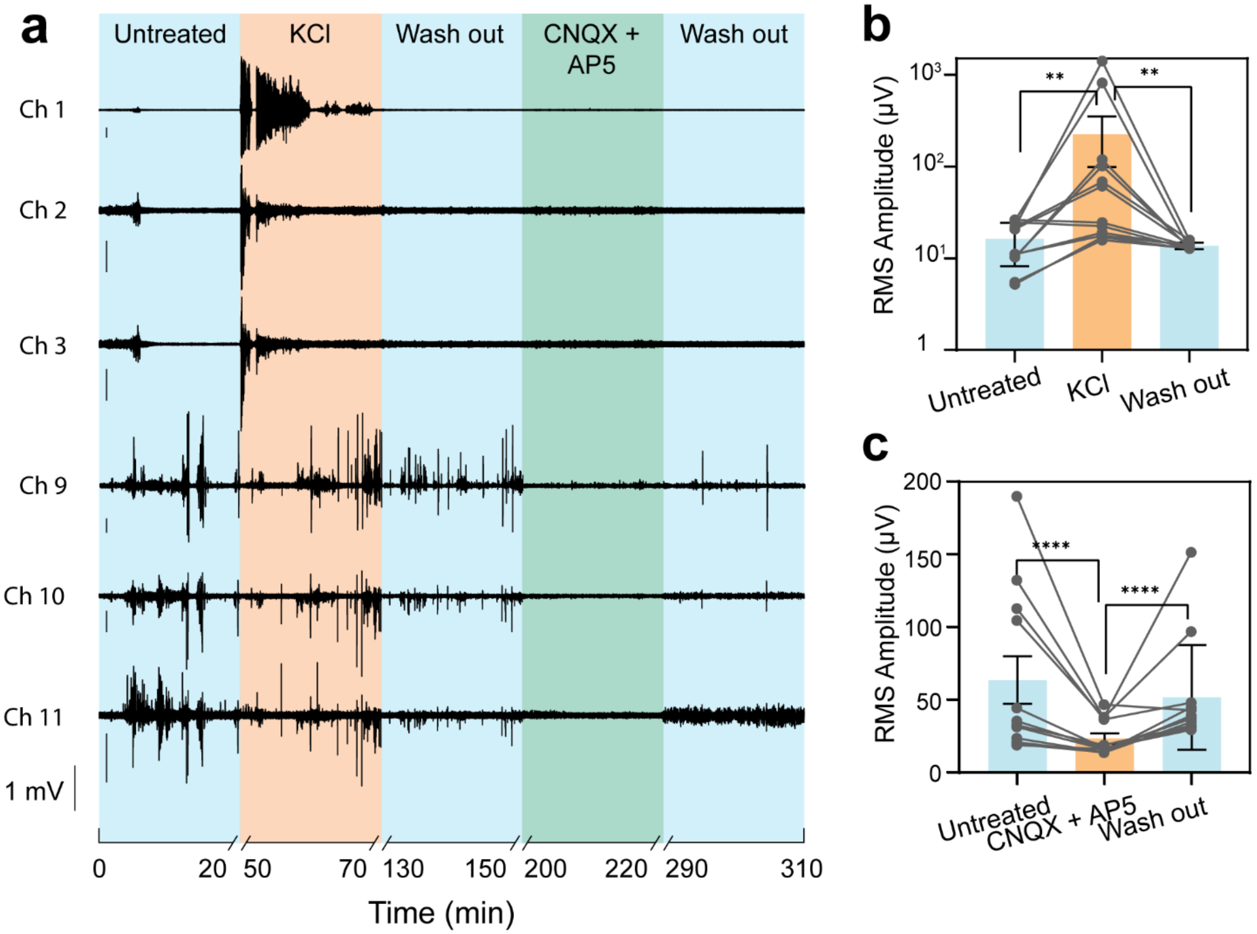


**Figure S10. Type A cyborg brain organoids drug response.** **a**, Raw voltage traces of 6 representative channels with a strong response to the injection of KCl or CNQX/D-AP5. **b**, Injection of KCl produces a significant increase in RMS amplitude of multiple electrodes. **c**, Injection of CNQX and D-AP5 produces a significant decrease in RMS amplitude of the same electrodes (n = 12, bar plots show mean ± S.D.). ** $P<0.01$, **** $P<0.001$ two-tailed, paired t-test.

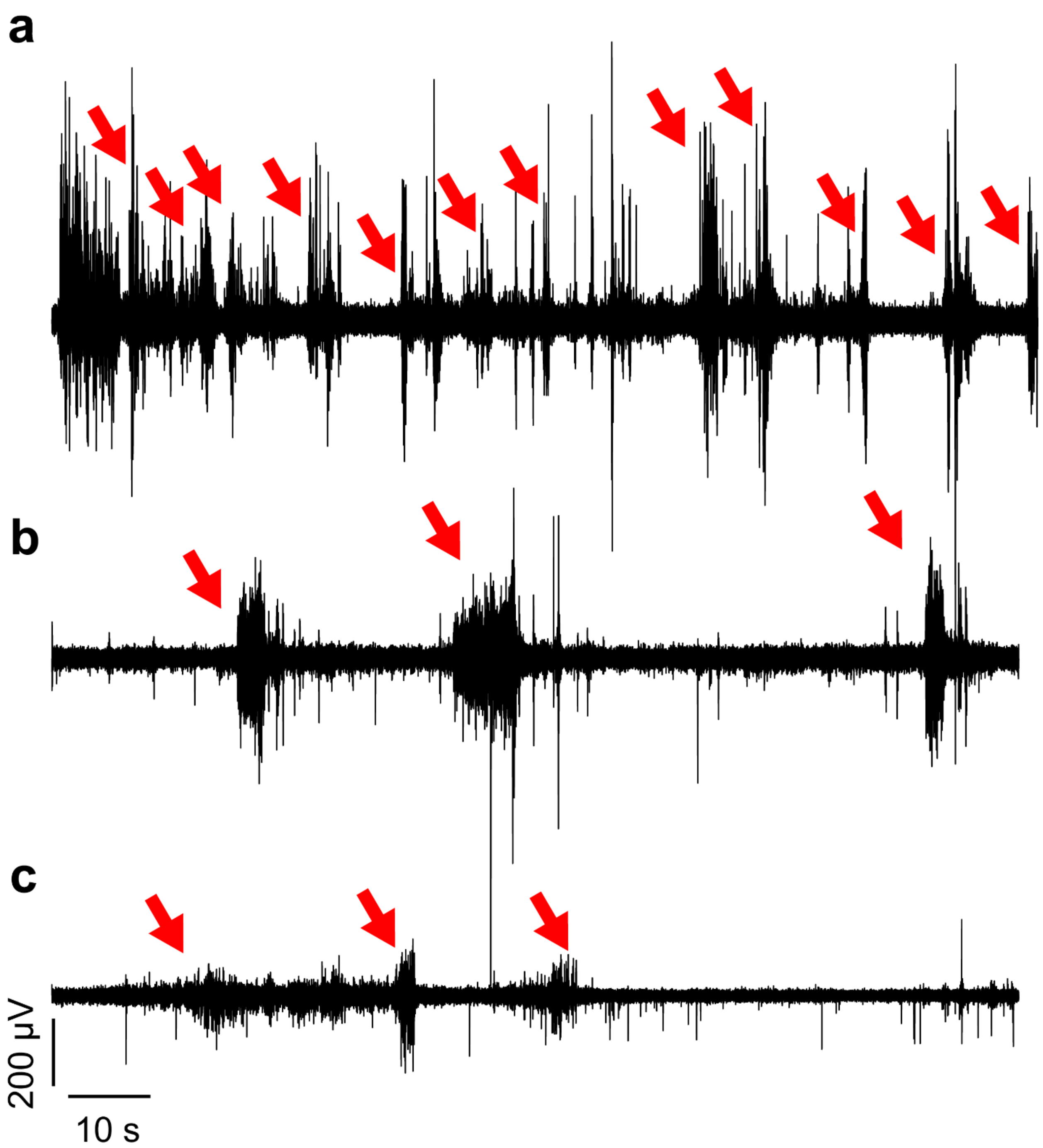


**Figure S11. Spontaneous bursting activity in type A cyborg brain organoids.** A 10 Hz high-pass filtered was applied to the signals. Red arrows show visible bursts. **a**, Organoid #5, channel 9. (**b**,**c**) Organoid #2, channel 9 (**b**) and 8 (**c**) (from different recordings).

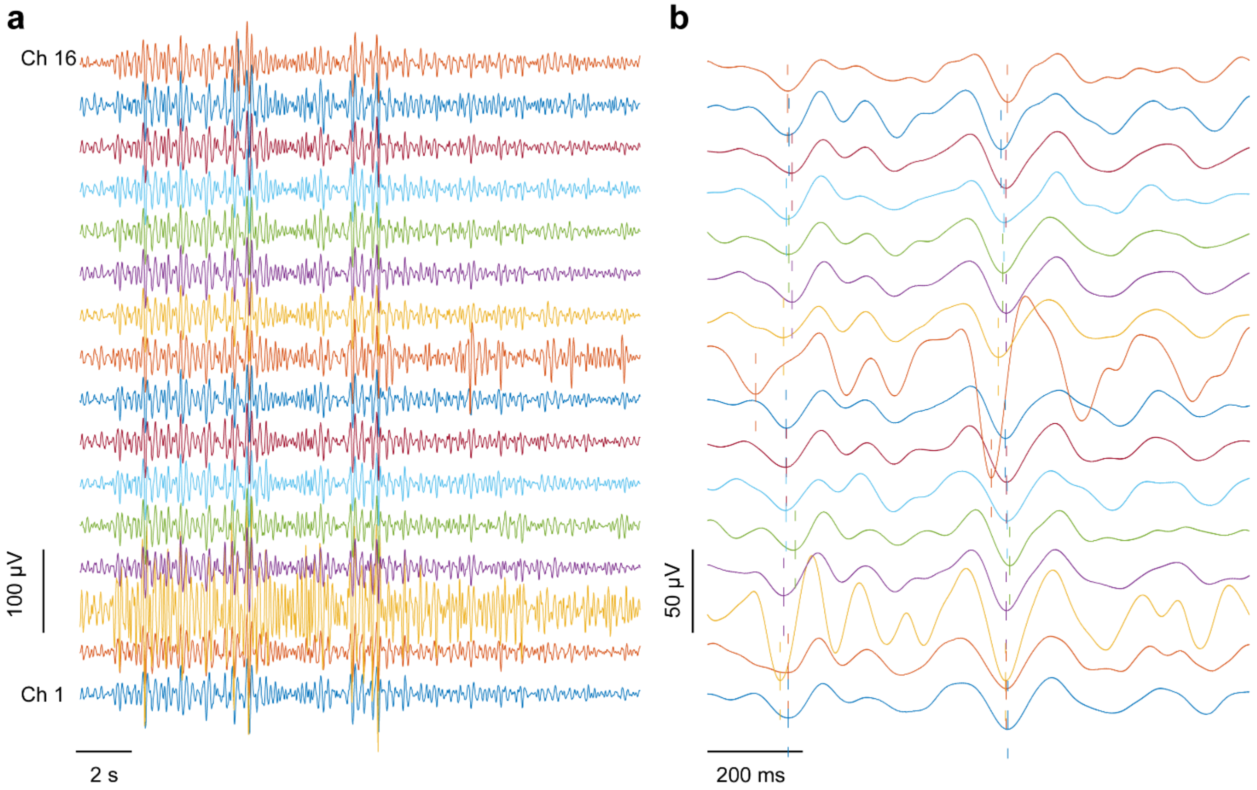


**Figure S12. Theta oscillations (4- 8 Hz bandpass) across all channels of the same device. a,** 20s window. **b**, Zoom-in of panel (**a**). Dashed lines help visualize the time delay of theta oscillations at a local minimum between different electrodes.

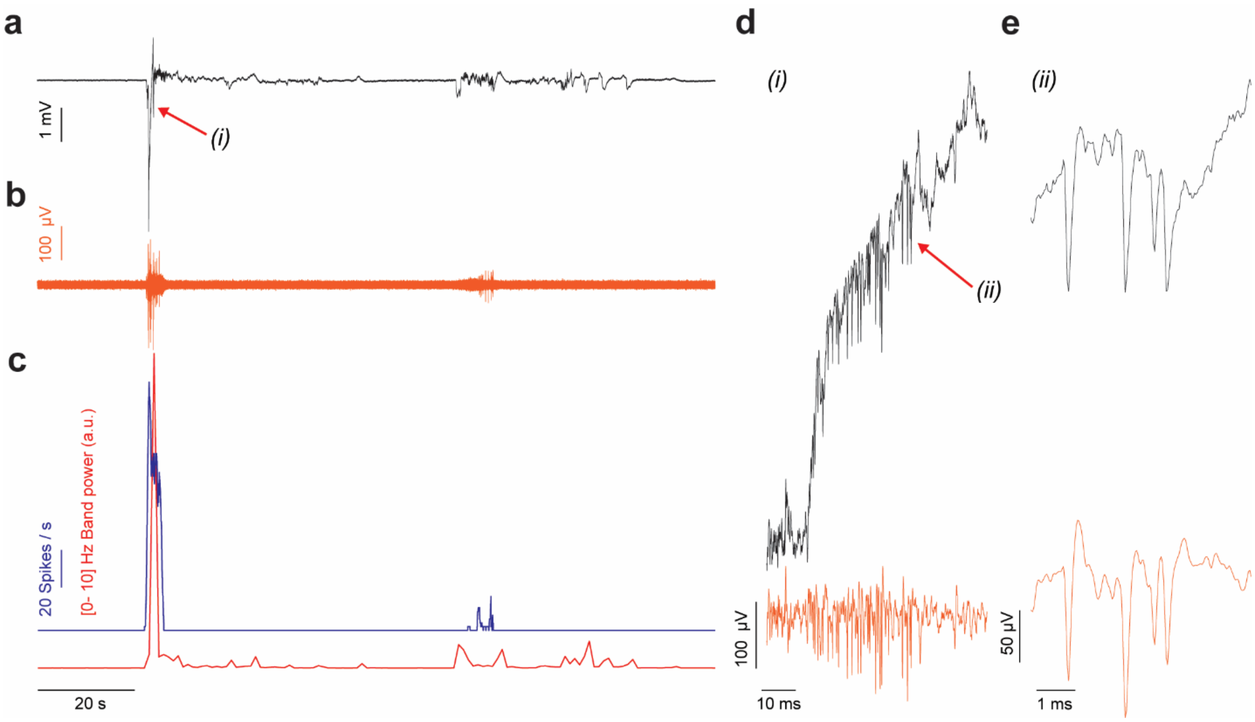


**Figure S13. LFP and spiking burst co-activation in a type B cyborg brain organoid. a**, raw voltage trace showing large amplitude LFP events. **b**, Filtered trace (300-6000 Hz). **c**, Moving average band power of the raw signal between 0 and 10 Hz (time bin of 1 s) is correlated with the moving average spiking rate (for all action potentials detected on the filtered trace of panel (**b**), with a time bin of 500 ms). **d**, Zoom-in of panels (**a, b**), around *(i)*. **e**, Zoom-in of panel (**d**), around *(ii)*.

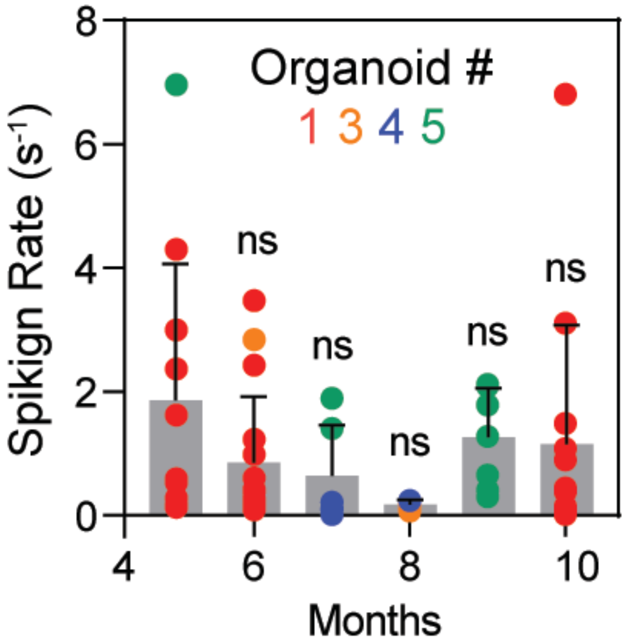


**Figure S14. Spiking rate of neurons detected in type A cyborg brain organoids.** One-way ANOVA with Month 5 data as control group shows no statistically significant change in spiking rate over time.

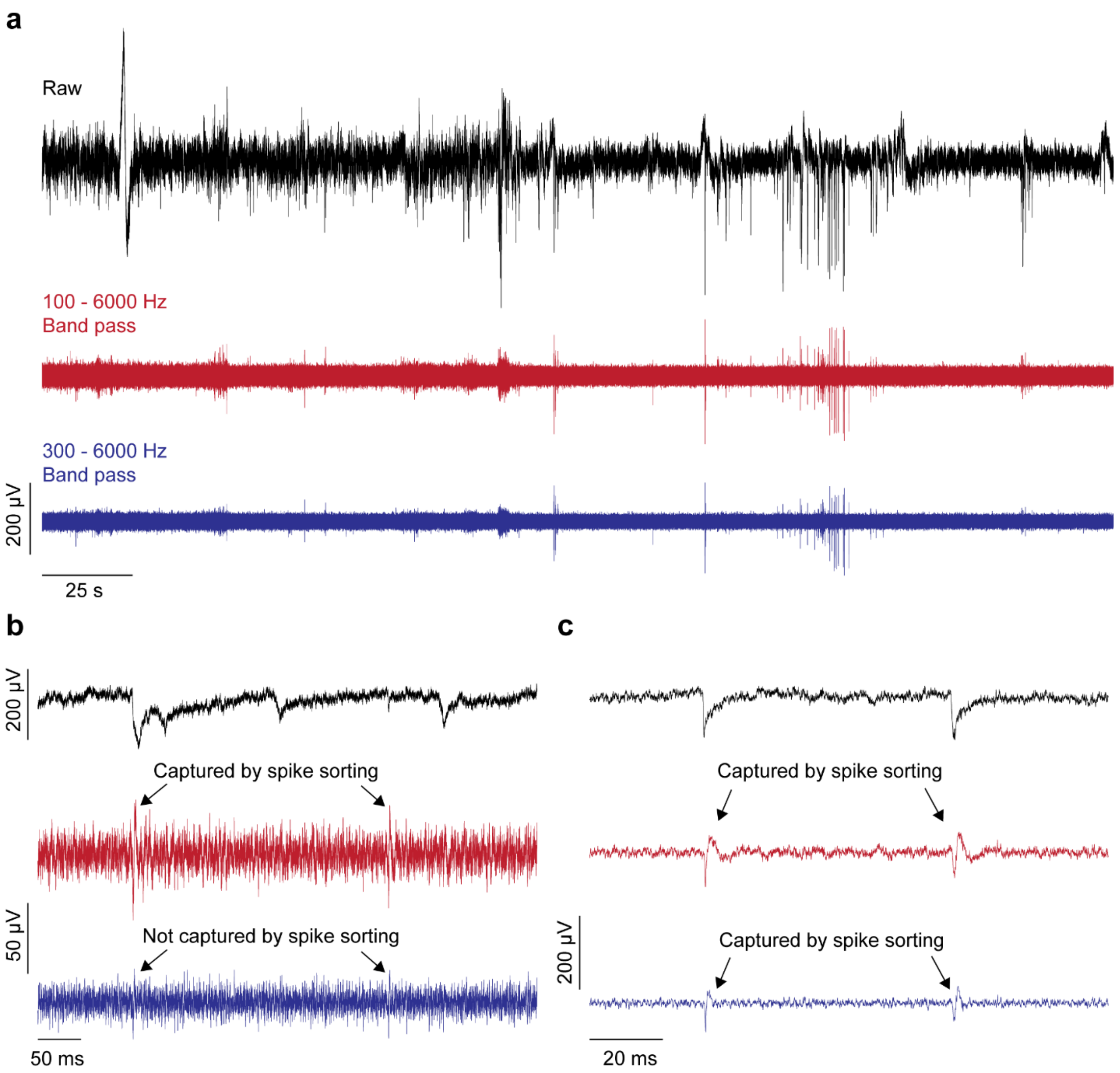


**Figure S15. Effect of 100 – 6000 Hz and 300 – 6000 Hz bandpass filters on spike detection. a**, Raw and filtered traces for a representative electrode in a type B organoid at month 2 post-assembly. **b**, Zoom-in of the voltage traces shows that slow spikes of the raw recording have a lower signal-to-noise ratio after 300 – 6000 Hz filtering (Bottom) than 100 – 6000 Hz filtering (Middle). Such spikes in the 300 – 6000 Hz filtered signal are not retained during spike detection. **c**, Zoom-in of the voltage traces shows that fast spikes of the raw recording have a sufficient signal-to-noise ratio after filtering to be retained during spike detection for both filters.

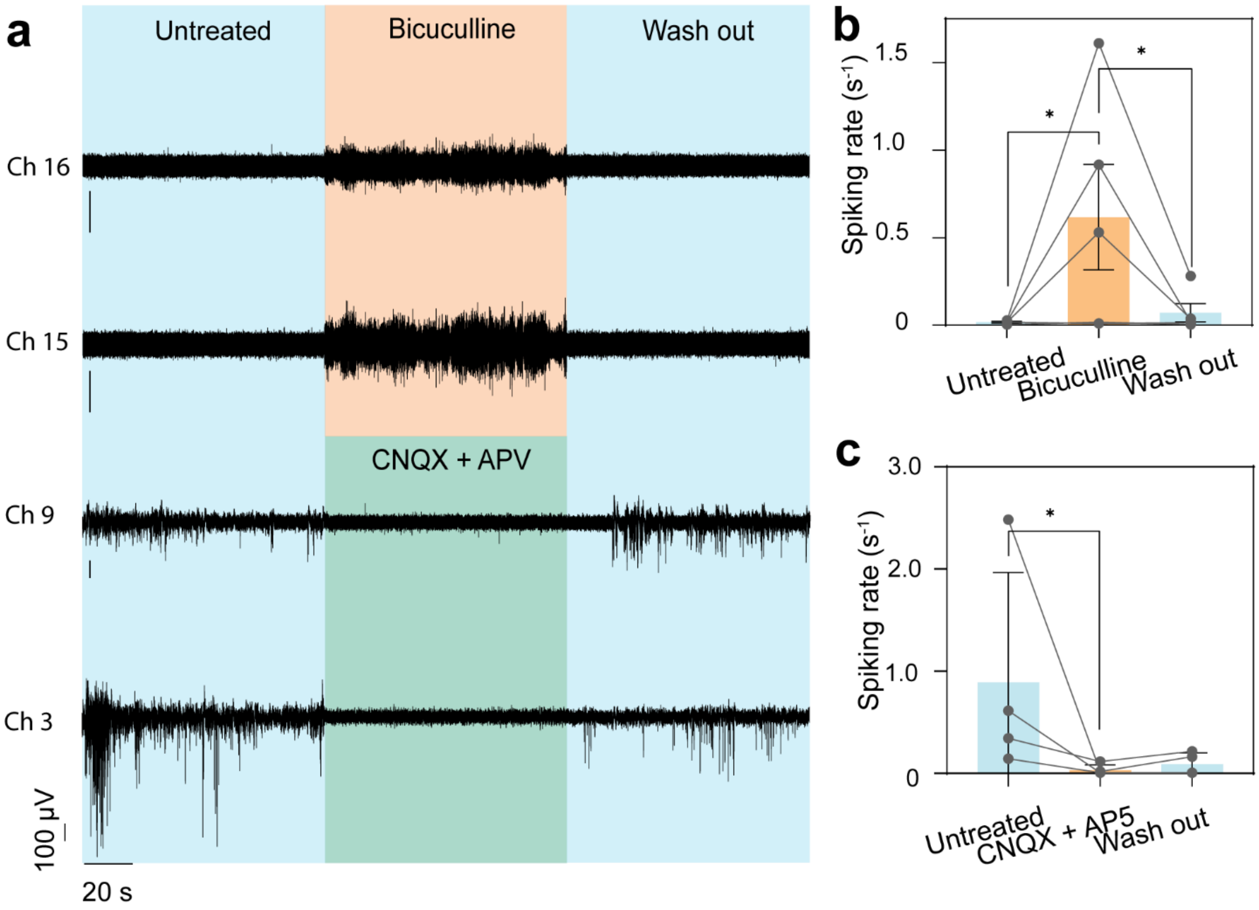


**Figure S16. Type B cyborg brain organoids drug response. a**, Filtered (300-3000 Hz Bandpass) voltage traces of 4 representative channels with a strong response to the injection of BCC or CNQX/D-AP5. **b**, Injection of BCC produces a significant increase in spiking rate of multiple electrodes. **c**, Injection of CNQX and D-AP5 produces a significant decrease in spiking rate of multiple electrodes (n = 4 electrodes including data from p = 2 different type B cyborg brain organoids, bar plots show mean ± S.D.). * $P<0.05$, two-tailed, paired t-test.

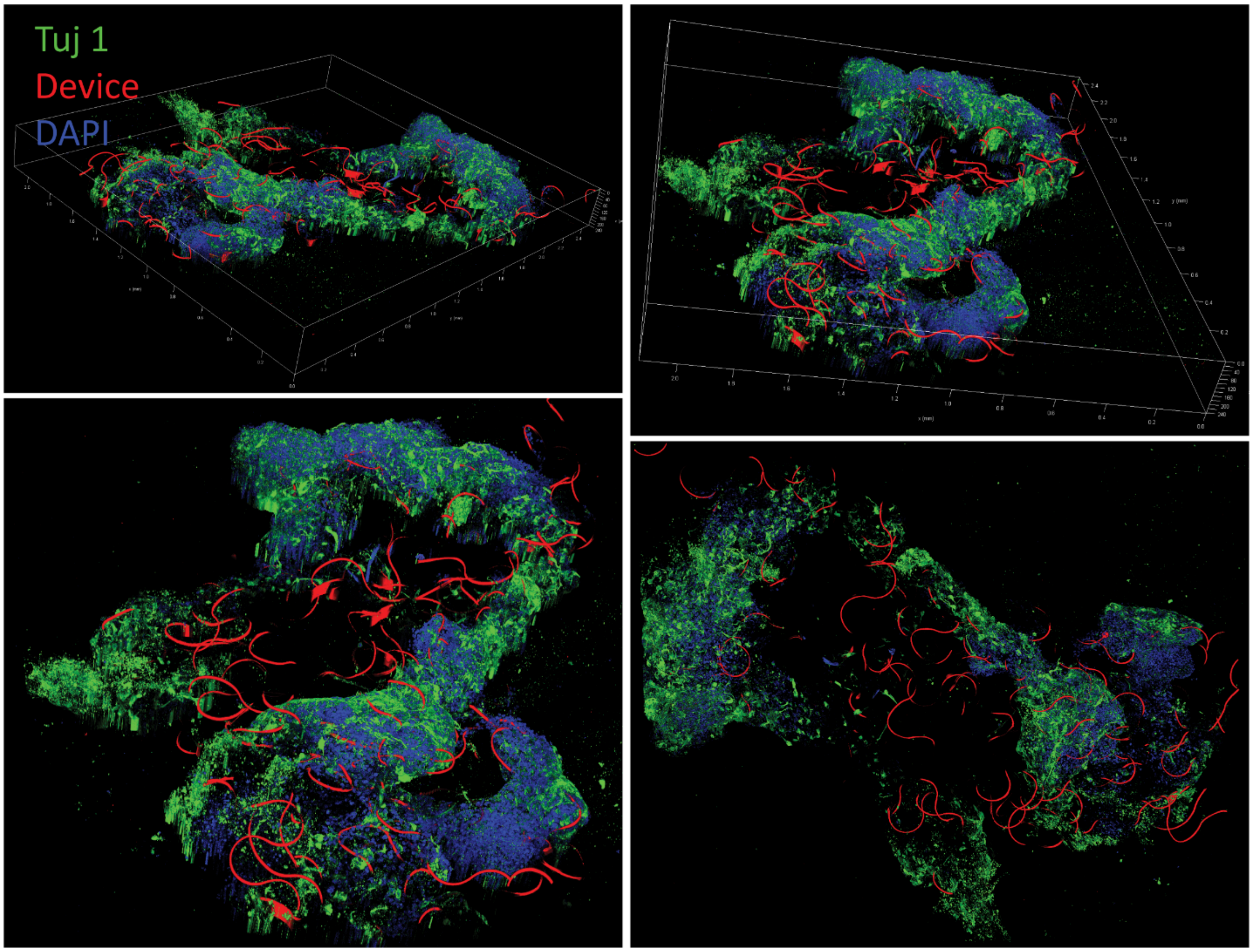


**Figure S17**. **3D reconstructed fluorescence images of a cleared, immunostained type B cyborg brain organoid at month 3 of differentiation.** Red, green and blue colors correspond to R6G, Tuj 1 and DAPI. The imaging parallelepiped has a volume of 2.4 x 2.2 x 0.24 $mm^3$.

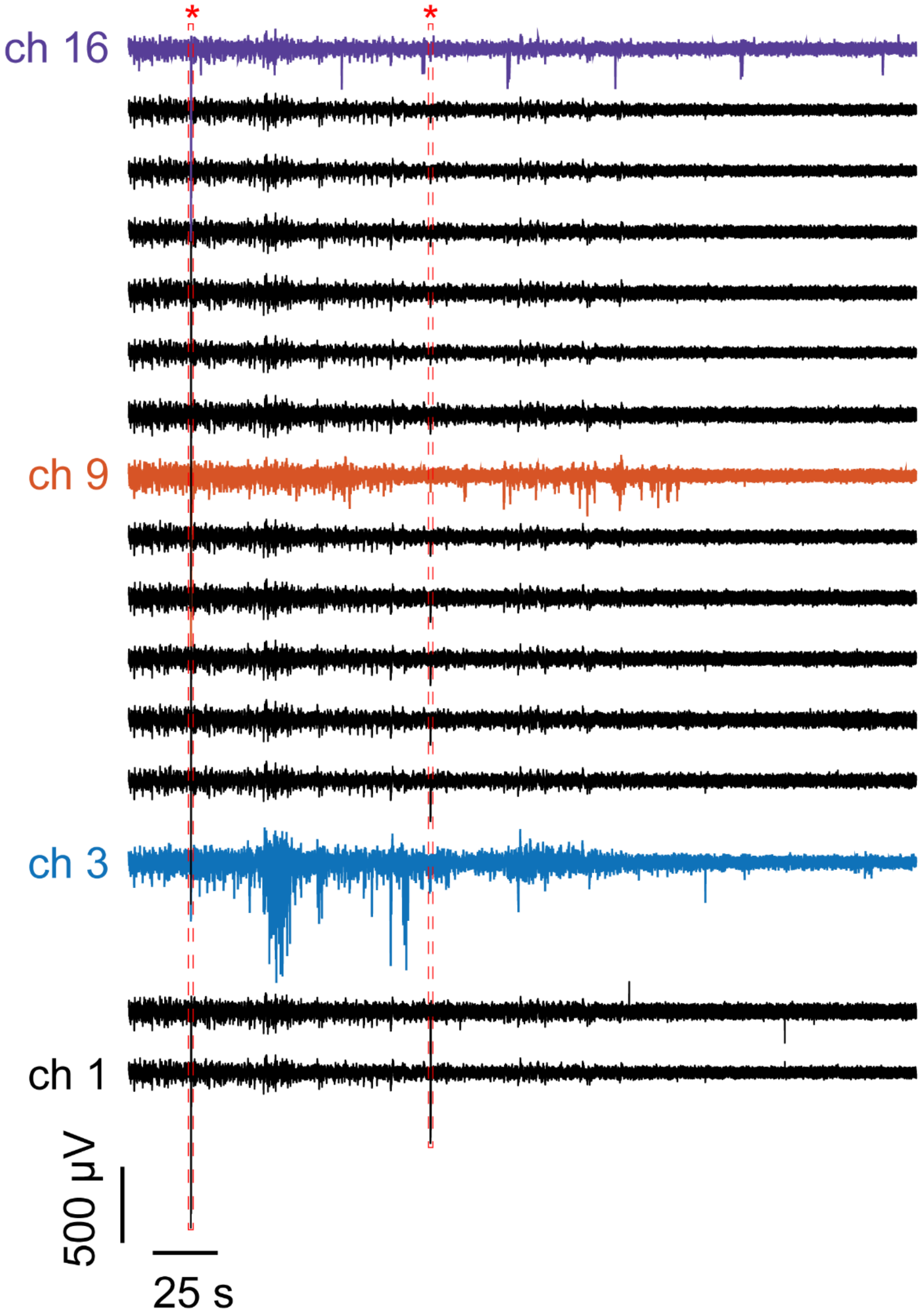


**Figure S18. Raw voltage recording of type B cyborg brain organoid.** Channel 3, 9 and 16 traces are plotted in Figure 5. Red asterisk and dashed border rectangles show voltage artefact common to all channels.

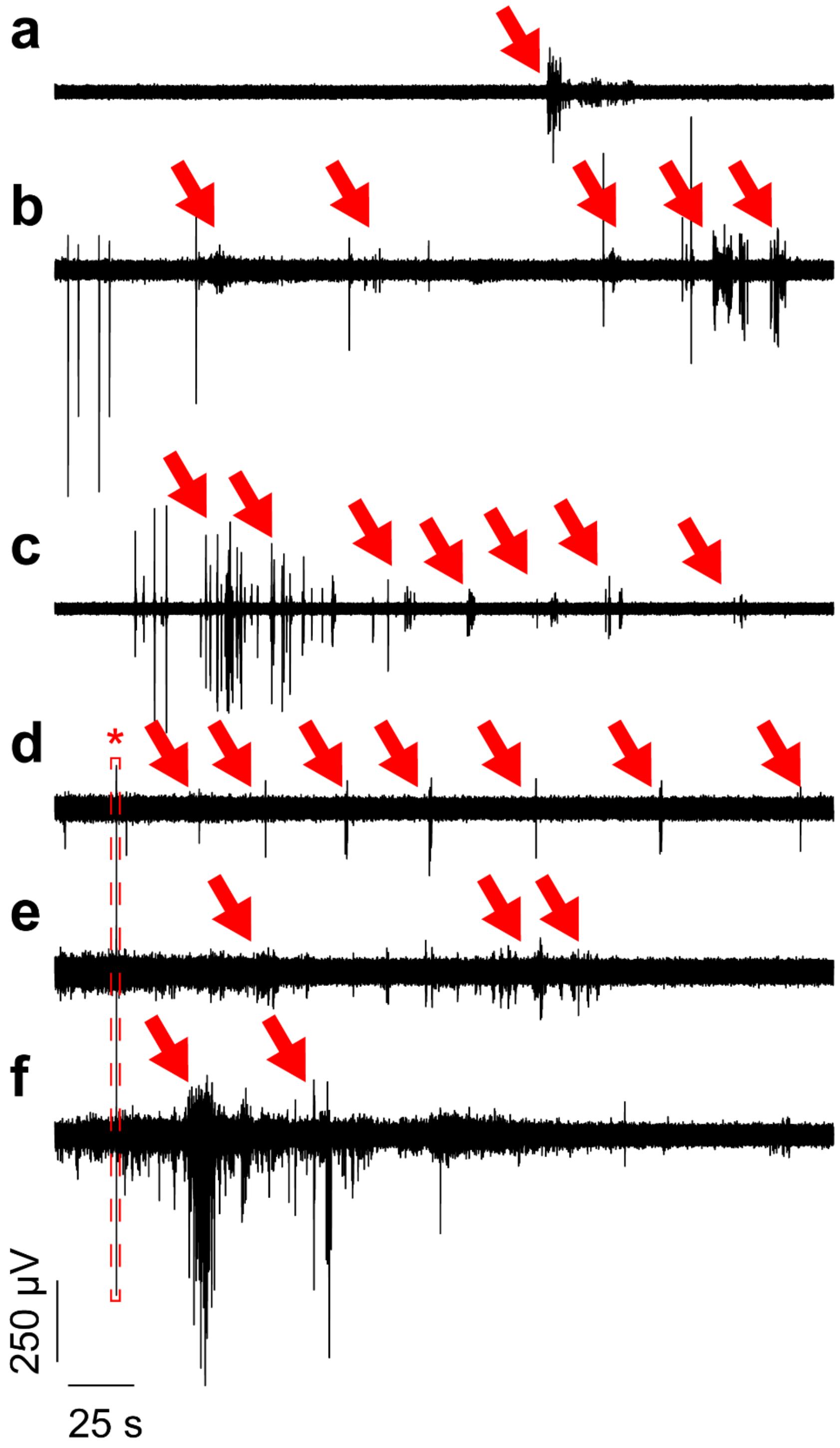


**Figure S19. Spontaneous bursting activity in type B cyborg brain organoids.** A 10 Hz high-pass filtered was applied to the signals. Red arrows show spiking bursts. Red asterisk and dashed border rectangles show voltage artefact common to all channels. **a**, Organoid #16, channel 15. **b**, Organoid #8, channel 11. **c**, Organoid #7, channel 8. **(d, e, f)** Organoid #6, channel 16 (**d**), 9 (**e**) and 3 (**f**) (from the same recording).
